# Comparative analysis of temperature preference behavior and effects of temperature on daily behavior in eleven *Drosophila* species

**DOI:** 10.1101/2022.04.18.488610

**Authors:** Fumihiro Ito, Takeshi Awasaki

**Affiliations:** Department of Biology, Kyorin University School of Medicine, 6-20-2 Shinkawa, Mitaka 181-8611 Tokyo, Japan

## Abstract

Temperature is one of the most critical environmental factors that influence various biological processes. Species distributed in different temperature regions are considered to have different optimal temperatures for daily life activities. However, how organisms have acquired various features to cope with particular temperature environments remains to be elucidated. In this study, we have systematically analyzed the temperature preference behavior and effects of temperatures on daily locomotor activity and sleep using 11 *Drosophila* species. We also investigated the function of antennae in the temperature preference behavior of these species. We found that, (1) an optimal temperature for daily locomotor activity and sleep of each species approximately matches with temperatures it frequently encounters in its habitat, (2) effects of temperature on locomotor activity and sleep are diverse among species, but each species maintains its daily activity and sleep pattern even at different temperatures, and (3) each species has a unique temperature preference behavior, and the contribution of antennae to this behavior is diverse among species. These results suggest that *Drosophila* species inhabiting different climatic environments have acquired species-specific temperature response systems according to their life strategies. This study provides fundamental information for understanding the mechanisms underlying their temperature adaptation and lifestyle diversification.

## Introduction

Temperature has been identified as one of the most critical environmental factors affecting various biological processes and species distribution [1]. Each species adapts to the temperature conditions of its habitat by tuning its lifestyle and life history. For instance, species living in higher latitudes or altitudes are known to tolerate cold and have evolved a higher capacity to maintain their daily life activities at lower temperatures [2-4]. However, the mechanism by which organisms have acquired such features remains unclear. Elucidating the mechanism of temperature adaptation and its diversification will significantly contribute to shed light on environmental adaptation and evolution.

The genus *Drosophila* includes about 1300 described species and is widely distributed from the tropical to subarctic regions [5, 6]. Several studies have explored their ecology and environmental adaptations such as food habits, temperature resistance, and diapause [7-12]. In particular, *D. melanogaster* is identified as a powerful and important model organism for biological research and has contributed to our understanding of the molecular and cellular mechanisms underlying various biological processes. Hence, *Drosophila* species are determined to be potentially useful in research in terms of for analyzing temperature adaptation and its diversification at molecular and cellular levels.

At a certain range of temperature, an animal exhibits active foraging and reproductive behaviors. In general, the temperature range at which animals remain active is dependent on the temperatures of their habitats [13]. For instance, animals living in cool climates generally exhibit higher activities even at low temperatures than animals living in tropical regions. However, animals are not always active; they can often change their activity level due to the effects of various factors even when ambient temperature is favorable. The activity level displays circadian rhythmicity. For example, *D. melanogaster* as well as several other animals exhibit circadian rhythmicity in locomotor activity; its inactive phase is often interpreted as “sleep” because it displays not only low locomotor activities but also low sensitivities to environmental signals [14]. Temperature has also been suggested to affect circadian rhythmicity; high temperature causes reduced daytime activity and increased nighttime activities [15], and total sleep length increases with increasing average temperature of habitats [16]. Nevertheless, there is limited research on the effects of temperatures on the locomotor activity and sleep of other *Drosophila* species from different climatic regions.

Ectothermic animals, whose body temperature is highly dependent on ambient temperature, require behavioral thermoregulation in response to temperature fluctuations. One of the behavioral thermoregulations is temperature preference behavior by which animals can maintain body temperature within a physiologically permissive range [17]. The genetic and cellular mechanisms underlying apparent temperature preference behavior in *D. melanogaster* have been extensively investigated [18-21]. Previous studies have shown that the transient receptor potential (TRP) ion channel A1 in the brain and Gustatory Receptor (GR) 28b in the antennae mediate warmth-sensing, and a trio of Ionotropic Receptors (IRs), IR21a, IR25a, and IR93a in the antennae mediate cooling [19, 22, 23]. Therefore, it is possible that the temperature preference of *D. melanogaster* is regulated by the balance between heat and cold or cooling avoidances through the activation of the TRP channel, GR and IRs. Hence, it would be useful to understand whether other *Drosophila* species have species-specific preferred temperatures and whether their preferred temperatures correlate with the optimal temperatures required for their activities and are determined by the thermal sensing system as in *D. melanogaster* [19, 22, 23].

Following the sequencing of the whole genome of *D. melanogaster*, the genome was also sequenced in other 11 *Drosophila* species that have been used for research in various fields to date [24]. This allows the analysis of the genetic mechanisms underlying unexplored biological phenomena that cannot be investigated by research using only *D. melanogaster*. The range of accessible research is broad, from morphology, physiology, and behavior to ecology and environmental adaptation mechanisms. Therefore, by analyzing how each *Drosophila* species adapts to its habitat temperatures, it is possible to understand a part of the mechanisms underlying temperature adaptation and its diversity. In this study, we have systematically analyzed the temperature preference behavior and effects of temperatures on daily locomotor activity and sleep in laboratory strains of 11 sequenced *Drosophila* species by applying the analysis systems developed in *D. melanogaster*. Furthermore, we analyzed the function of antennae, which is indispensable for temperature sensing in *D. melanogaster*, in the temperature preference behavior of these species. Our results demonstrate that the effects of temperature on daily activities and thermoregulatory behavior have been diverse among *Drosophila* species and that *Drosophila* species inhabiting unique climatic environments have acquired species-specific temperature response systems based on their own life strategies and evolutionary processes.

## Results

### Effects of temperature on total daily locomotor activities

To understand the effect of temperature on the daily behavior of *Drosophila* species distributed in different temperature regions, we examined the daily locomotor activity at different temperatures in the following 11 sequenced *Drosophila* species: cosmopolitan (*D. melanogaster* and *D. simulans*), tropical (*D. ananassae, D. erecta, D. yakuba*, and *D. sechellia*), subtropical (*D. willistoni* and *D. mojavensis*), and temperate (*D. persimilis, D. pseudoobscura*, and *D. virilis*) species. Using the *Drosophila* Activity Monitor system [25], we were able to analyze the amount of daily locomotor activity quantitatively at five experimental temperatures, i.e., 17°C, 20°C, 23°C, 26°C, and 29°C. As the viability of the adults of *D. persimilis* and *D. pseudoobscura* was low at 29°C, these two species were analyzed at only four experimental temperatures. First, we compared the amount of daily locomotor activities among these *Drosophila* species (Supplementary Fig. 1). The ranges of the total daily activity were quite diverse in these species (Kruskal–Wallis test: χ^2^ = 833.18, *p* < 0.001). This analysis revealed that among these species, *D. melanogaster* is most active, whereas *D. erecta* and *D. mojavensis* are less active. The average amount of daily locomotor activity in *D. melanogaster* was ten times higher than that of *D. erecta* and *D. mojavensis*. This result suggested that each *Drosophila* species has its own range of total daily locomotor activity.

Next, we examined the effect of temperature on the amount of daily locomotor activity (Fig. 1). In *D. melanogaster*, temperature was found to have no significant effect on the amount of activity (Kruskal–Wallis test: *D. melanogaster*: χ^2^ = 8.49, *p* = 0.08), implying that *D. melanogaster* maintains its activity levels almost constantly in the wide range of temperatures. Although there was no significant difference in the amount of activity from 17°C to 26°C in *D. pseudoobscura*, this species died at 29°C in a few days, suggesting that *D. pseudoobscura* finally loses its locomotor activity at 29°C. The other nine species exhibited significant differences in the amount of activity among the experimental temperatures (Kruskal–Wallis test: *p* < 0.05). The amount of activity was higher at ≥ 26°C than at < 26°C in *D. sechellia* and *D. mojavensis*. In *D. ananassae, D. yakuba*, and *D. willistoni*, the activity level was significantly higher at ≥ 23°C than at < 23°C. However, in *D. simulans* and *D. persimilis*, the amount of activity was generally higher at ≤ 23°C than at > 23°C. In *D. erecta*, the peak temperature for the amount of activity was 23°C. In *D. virilis*, the amount of activity was higher at ≤ 20°C than at > 20°C. To estimate the optimal temperature and maximum performance for daily locomotor activity, we fitted thermal performance curve (TPC) model for each species (Supplementary Fig. 2). These results, in addition to the effects of temperature on the amount of activity and optimal temperatures estimated by TPC models (with some exceptions), indicate that tropical and temperate *Drosophila* species tend to exhibit higher activity levels at high and low temperatures, respectively.

**Figure 1.**
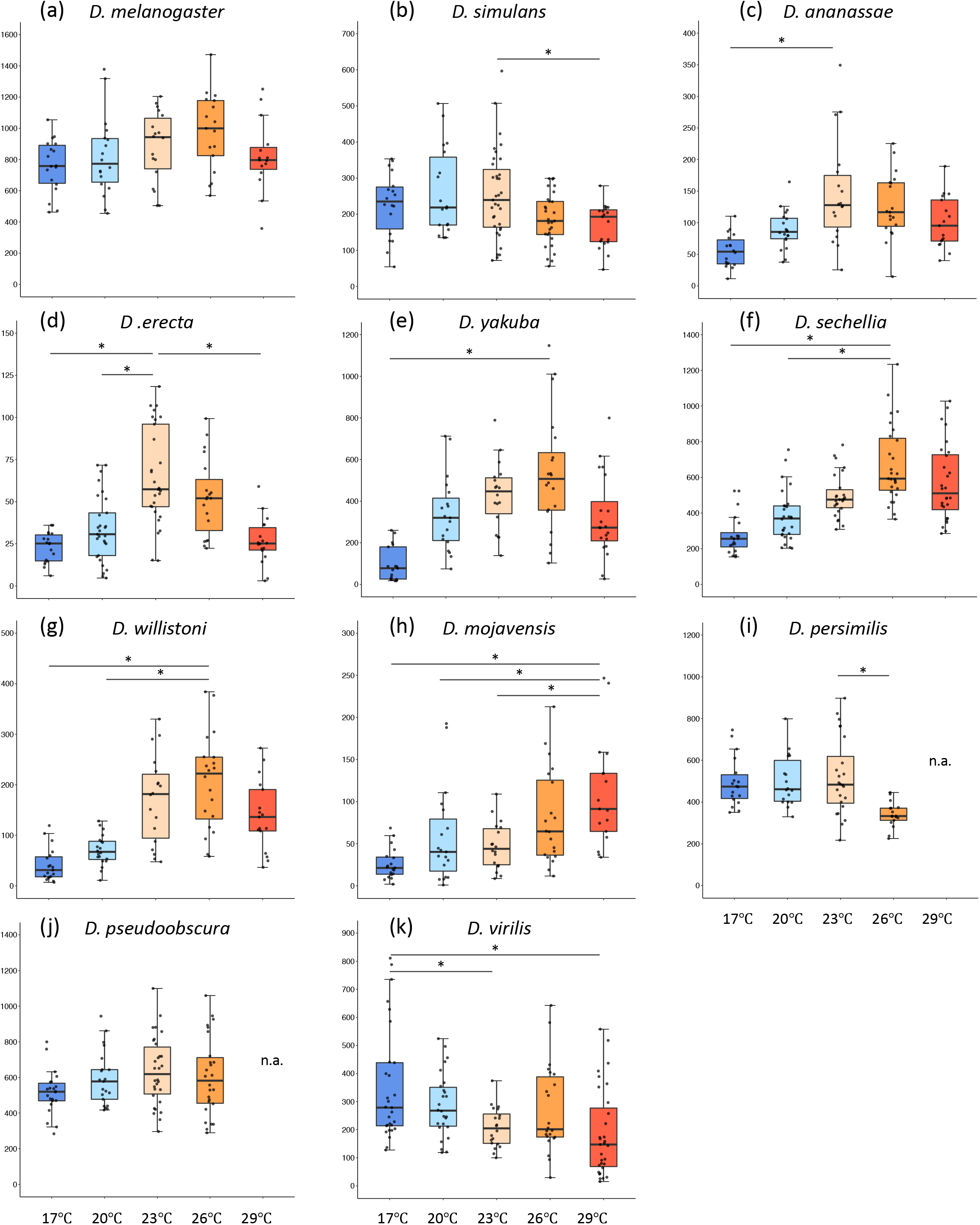
Total daily locomotor activity of 11 *Drosophila* species at five different temperatures. (a) ∼ (k) represents the total daily locomotor activity in each species; (a) *D. melanogaster*, (b) *D. simulans*, (c) *D. ananassae*, (d) *D. erecta*, (e) *D. yakuba*, (f) *D. sechellia*, (g) *D. willistoni*, (h) *D. mojavensis*, (i) *D. persimilis*, (j) *D. pseudoobscura*, and (k) *D. virilis*. The y axis shows the activity levels (counts) and x axis shows time of the day. Caution that the y axis range are different among species. The box color indicates experimental temperatures; dark blue:17°C, light blue: 20°C, light orange: 23°C, dark orange: 26°C, red: 29°C. Except for *D. melanogaster* and *D. pseudoobscura*, all the other species show the significant difference in total locomotor activity among different temperature conditions (Kruskal–Wallis tests, *p* < 0.05). In this and following figures, each box plot shows the median as the bold line, first and third quartiles as the box boundaries, and 1.5 times of the interquartile ranges as whiskers. Each dot indicates average total daily activity (counts) of individual fly. The horizonal bar with * above the boxes indicates the significant differences in the pairwise-comparison between the highest average of total locomotor activity and others (Bonferroni/Dunn test, *p* < 0.05). The sample size of each analysis is shown in Supplementary Fig. 2.

### Effects of temperature on the daily activity pattern

We examined how temperature affects the daily activity pattern in these *Drosophila* species. The daily patterns of locomotor activity were investigated at different temperatures (Fig. 2, Supplementary Fig. 3). Each *Drosophila* species exhibited a unique daily activity pattern. Most species have bimodal morning and evening peaks of activity in a day, as also known in *D. melanogaster* [26]. However, the height, shape, and ratio of the two peaks were noted to vary among species. To investigate the effect of temperature on activity peaks, we focused on the amount of activity in the first (from 2:00 to 14:00) and second (14:00–2:00) half of a day, which includes the morning and evening peak, respectively (Supplementary Fig. 4). In all the examined species, either or both the amounts of locomotor activity in the first and second half of the day were affected by temperature (Kruskal–Wallis test: *p* < 0.05, Supplementary Fig. 4). However, the ratio of the activity level in the first and second half of the day was almost constant at all experimental temperatures in most species (Supplementary Fig. 5). In fact, the overall activity peaks, for example, higher morning peak in *D. yakuba* and higher evening peak in *D. simulans*, were maintained in most species (Fig. 2 and Supplementary Fig. 3). Remarkably, although the ratio of activity was different between low and high temperatures in *D. willistoni*, this was caused by the disappearance of the morning and evening peaks at low temperature (≤ 20°C) rather than an inversion of the ratio. These results illustrated that the species-specific ratio of the morning and evening activity peaks is less affected by temperature, suggesting that each species maintains daily activity pattern even at different temperatures.

**Figure 2.**
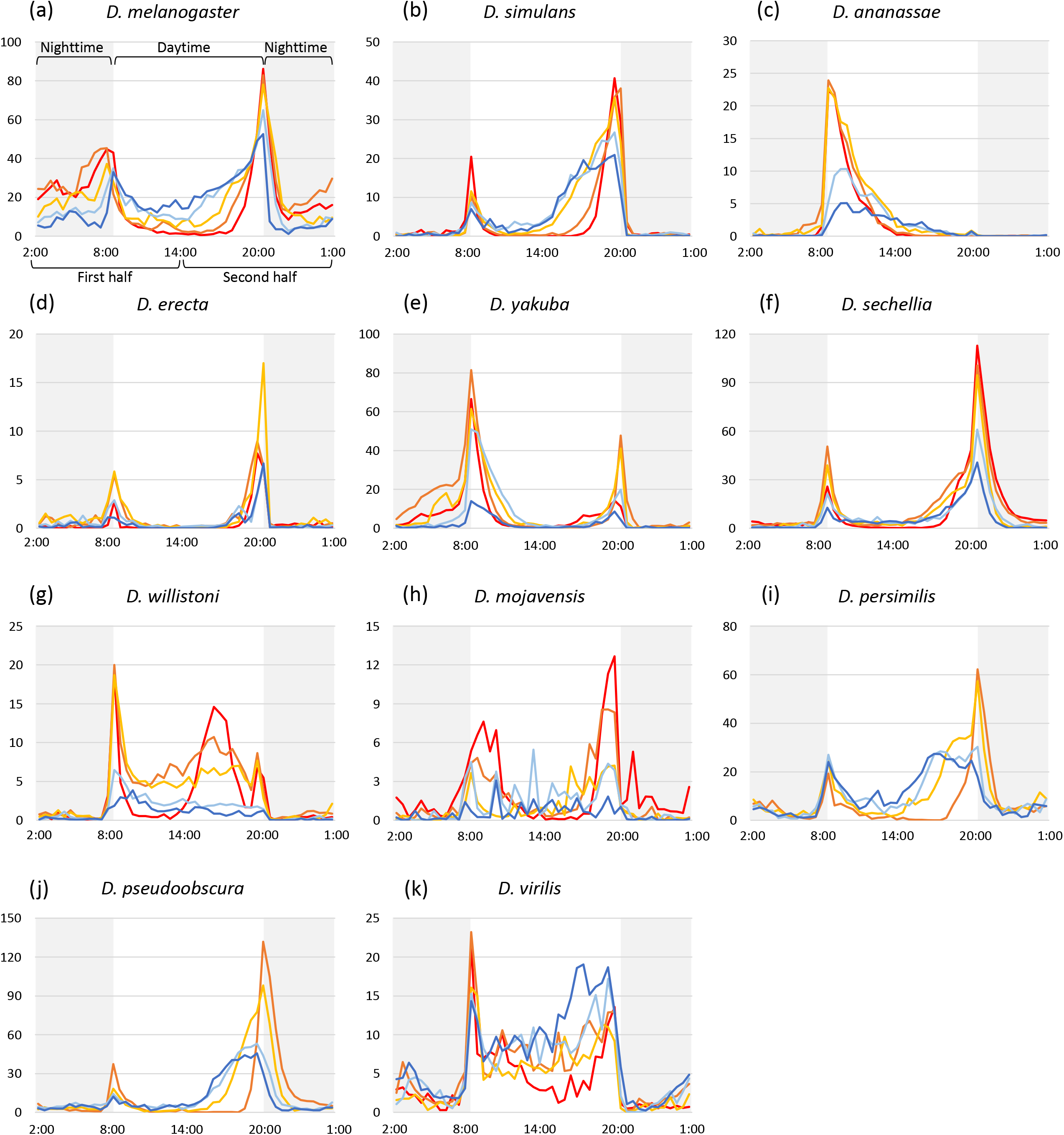
Profiles of daily locomotor activity in 11 *Drosophila* species at five different temperatures. (a) ∼ (k) represents the profiles of daily locomotor activities in each species; (a) *D. melanogaster*, (b) *D. simulans*, (c) *D. ananassae*, (d) *D. erecta*, (e) *D. yakuba*, (f) *D. sechellia*, (g) *D. willistoni*, (h) *D. mojavensis*, (i) *D. persimilis*, (j) *D. pseudoobscura*, and (k) *D. virilis*. The y axis shows the activity levels (counts). Caution that the y axis range are different among species. The lines depict the average counts in 30 min at each temperature condition. The line color indicates experimental temperatures; dark blue:17°C, light blue: 20°C, light orange: 23°C, dark orange: 26°C, red: 29°C. The shadow in the figure indicates the dark condition (nighttime).

To further explore the effect of temperature on the daily activity pattern, we have examined how light condition affect daily activity patterns (Fig. 3). All the examined species showed significant differences in both daytime (light on) and nighttime (light off) activity levels among the experimental temperatures (Kruskal–Wallis test: *p* < 0.05). At all experimental temperatures, the daytime activity level was higher or equal to the nighttime activity level in six species (daytime-active species), *D. simulans, D. ananassae, D. yakuba, D. willistoni, D. mojavensis*, and *D. virilis*. In *D. erecta*, the activity level scarcely differed between day- and nighttime. In contrast, nighttime activities were low at low temperatures, gradually increased with increasing temperature, and finally exceeded daytime activities at 23°C, 26°C, or 29°C in the remaining four species, *D. melanogaster, D. sechellia, D. persimilis*, and *D. pseudoobscura*. Regarding daytime/nighttime activity ratio, we observed three different patterns in the effect of temperature (Supplementary Fig. 6) as follows: (1) the daytime/nighttime ratio was not affected by temperature (*D. erecta, D. willistoni*, and *D. virilis*), (2) the ratio constantly decreased with increasing temperature (*D. melanogaster, D. simulans, D. sechellia, D. persimilis*, and *D. pseudoobscura*), and (3) the ratio was affected by temperature without a constant tendency (*D. ananassae, D. yakuba*, and *D. mojavensis*). These results indicate that some species evenly change their daytime and nighttime activity level depending on the temperature, while the others change the ratio of the daytime and nighttime activity level depending on the temperature.

**Figure 3.**
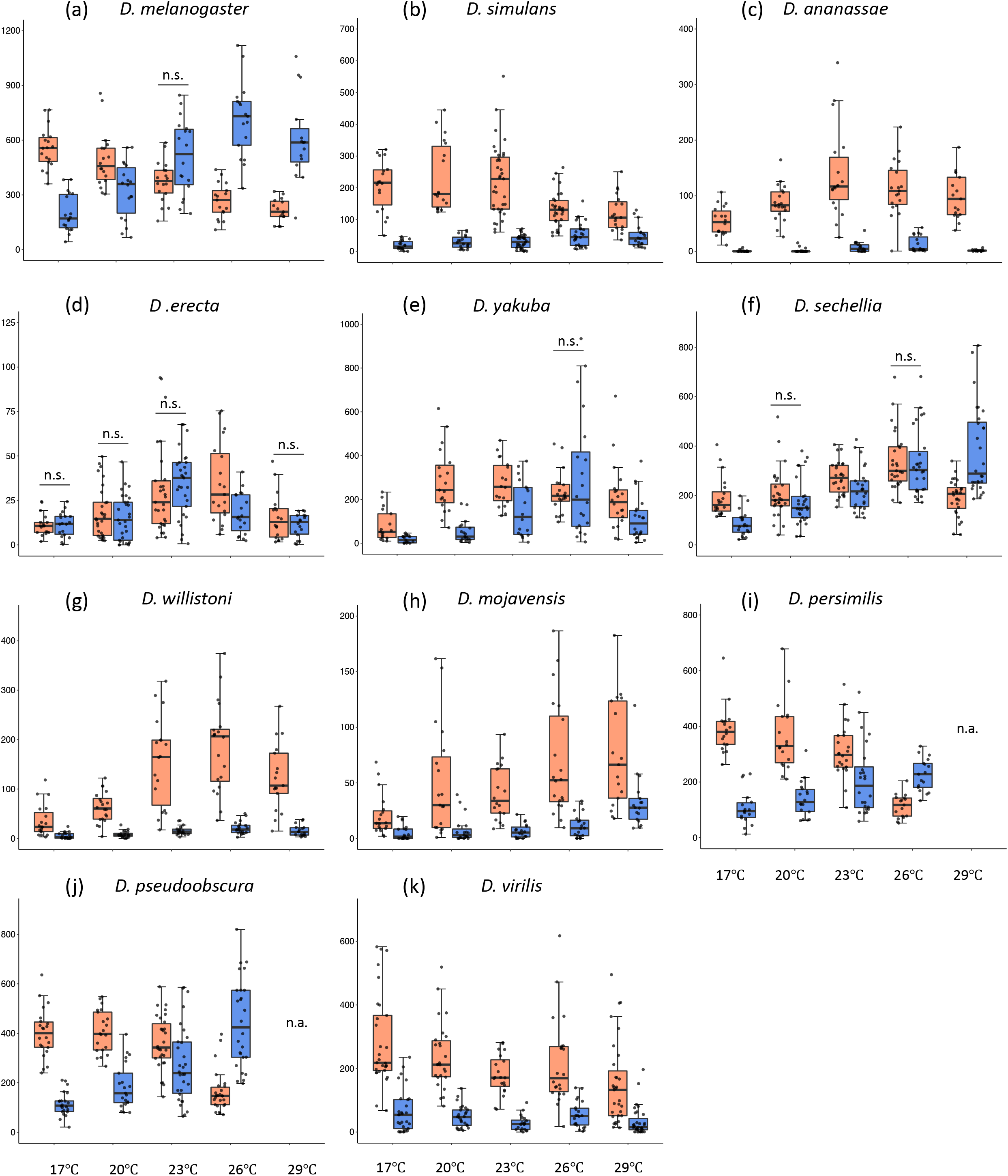
Daytime and nighttime locomotor activity of 11 *Drosophila* species at five different temperatures. (a) ∼ (k) represents the daytime and nighttime locomotor activity in each species; (a) *D. melanogaster*, (b) *D. simulans*, (c) *D. ananassae*, (d) *D. erecta*, (e) *D. yakuba*, (f) *D. sechellia*, (g) *D. willistoni*, (h) *D. mojavensis*, (i) *D. persimilis*, (j) *D. pseudoobscura*, and (k) *D. virilis*. The y axis shows the activity levels (counts). Caution that the y axis range are different among species. Box plots are shown as Figure 2. Orange box indicates daytime and blue box indicates nighttime locomotor activities. The horizonal bars with n.s. above the box daytime and nighttime pairs indicates not significant differences (Mann–Whitney U tests, *p* > 0.05), otherwise each pairs shows a significant difference (Mann–Whitney U tests, *p* < 0.05). In all species, the amount of daytime and nighttime activities is significantly affected by temperatures (Kruskal–Wallis tests, *p* < 0.05).

### Effects of temperature on sleep

Environmental temperature is known to influence sleep in several animals, including *D. melanogaster* [27-28]. However, how temperature affects sleep in other *Drosophila* species has not been well investigated. Thus, in this study, we examined the effect of temperature on sleep in the different *Drosophila* species (Fig. 4 and Supplementary Fig. 7). First, we compared the range of total length of sleep in a day and found that the ranges of daily sleep length were quite different (Kruskal–Wallis test: χ^2^ = 747.72, *p* < 0.001, Supplementary Fig. 8). This result suggests that, as observed in the locomotor activity, each species has its own range of daily sleep length, and this range is diverse among *Drosophila* species. The daily activity level and sleep length showed a significant negative correlation (Spearman’s rank correlation = -0.936, *p* < 0.05), indicating that long-sleep species are less active and short-sleep species are more active.

**Figure 4.**
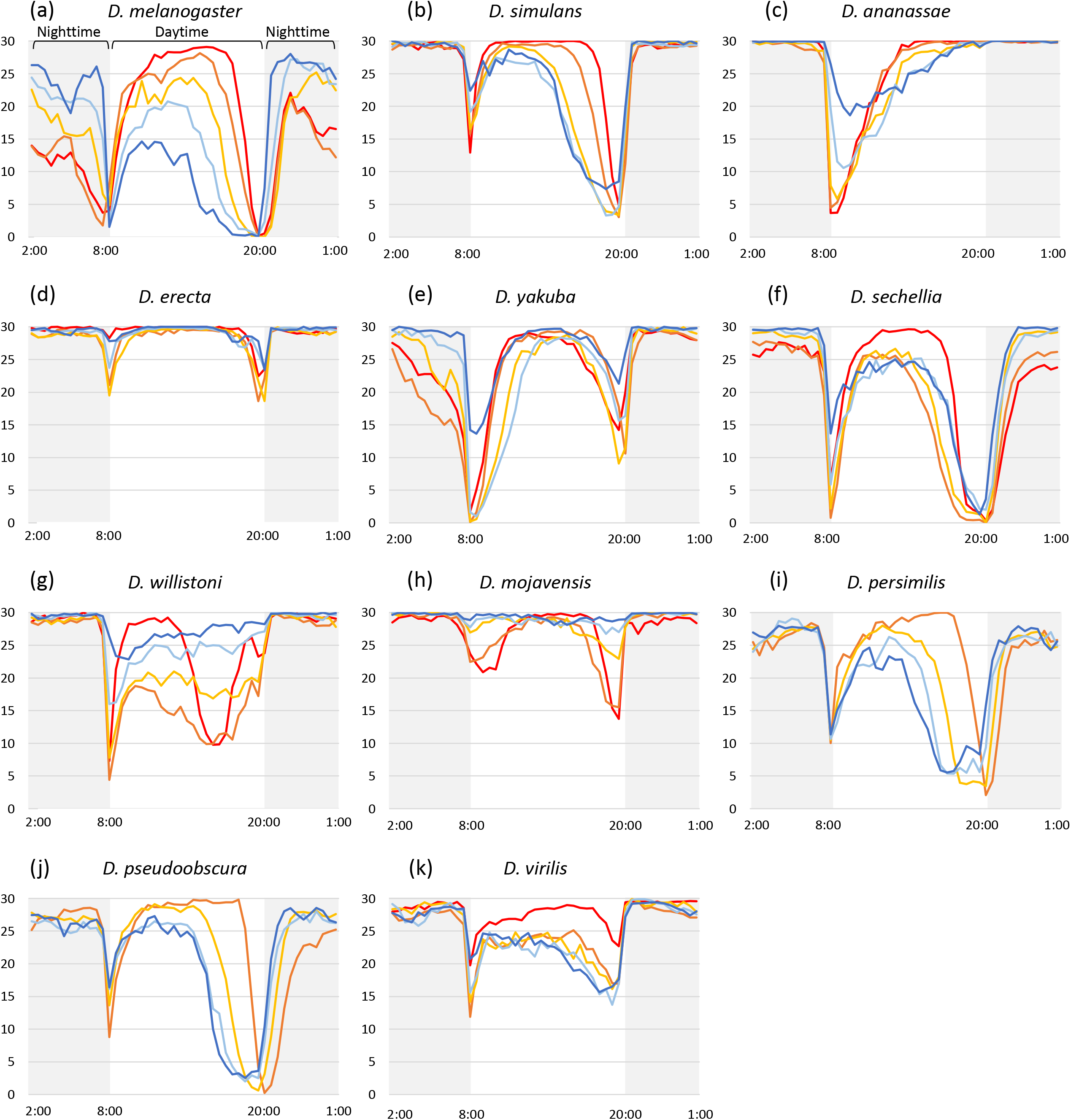
Profiles of daily sleep in 11 *Drosophila* species at five different temperatures. (a) ∼ (k) represents the daily sleep profile in each species; (a) *D. melanogaster*, (b) *D. simulans*, (c) *D. ananassae*, (d) *D. erecta*, (e) *D. yakuba*, (f) *D. sechellia*, (g) *D. willistoni*, (h) *D. mojavensis*, (i) *D. persimilis*, (j) *D. pseudoobscura*, and (k) *D. virilis*. The y axis shows the duration of sleep in 30 min. The lines depict the average of sleep duration in 30 min at each temperature condition. The line color indicates experimental temperatures; dark blue:17°C, light blue: 20°C, light orange: 23°C, dark orange: 26°C, red: 29°C. The shadow in the figure indicates the dark condition (nighttime).

Next, we compared the daily sleep length and patterns among different temperatures (Fig. 4 and Supplementary Fig. 9). *D. melanogaster* showed no significant difference in sleep length among different temperatures (Kruskal–Wallis test: χ^2^ = 2.33, *p* = 0.68), whereas the sleep length of all the other species was significantly affected by temperature (Kruskal–Wallis test: *p* < 0.05). In individual species, the sleep length and locomotor activity also showed negative correlation at different temperatures (Supplementary Table 1). Regarding the daily sleep pattern, all the examined species showed an opposite sleep pattern to the activity pattern (comparison between Fig. 2 and Fig. 4). Hence, the timing of increased and decreased sleep was almost constant at all the examined temperatures in all species. Moreover, the total day- and nighttime sleep was significantly affected by temperature in all species (Kruskal–Wallis test: *p* < 0.05, Supplementary Figs. 10 and 11).

We have also examined their length of single sleep and number of sleep in a day, which are the other aspects of sleep behavior (Supplementary Figs. 12 and 13). Excluding *D. melanogaster*, the single sleep duration and number of sleep were significantly affected by temperature in these species (Kruskal–Wallis test: *p* < 0.05). We observed that the total sleep length correlated positively with the length of single sleep and negatively with the number of sleep (Supplementary Table 1), suggesting that flies increase the total sleep length by increasing the length of single sleep and not by increasing the frequency of sleep.

### Temperature preference behavior

We next investigated the behavioral thermoregulation in these *Drosophila* species. First, we examined the distribution of flies on the temperature gradient field (Fig. 5). Each species exhibited species-specific distribution on the field. Of the 11 species, 7 showed prominent distribution peaks on the field, while *D. melanogaster* showed a distribution peak at 25°C as reported in a previous study [19, 21]. *D. simulans, D. erecta, D. persimilis*, and *D. pseudoobscura* showed peaks at 23°C, whereas *D. mojavensis* and *D. virilis* showed peaks at the higher temperatures of 25°C and 29°C, respectively. Although these seven species exhibited prominent distribution peaks, the shapes of the peak varied among them. *D. melanogaster* and *D. erecta* showed high and sharp peaks, whereas the other five species showed lower and gentle peaks. Among the remaining four species, *D. yakuba* and *D. sechellia* showed broad distributions at ≤ 23°C or ≤ 25°C, respectively, with no apparent peaks. *D. ananassae* and *D. willistoni* swarmed on the colder side of the field.

**Figure 5.**
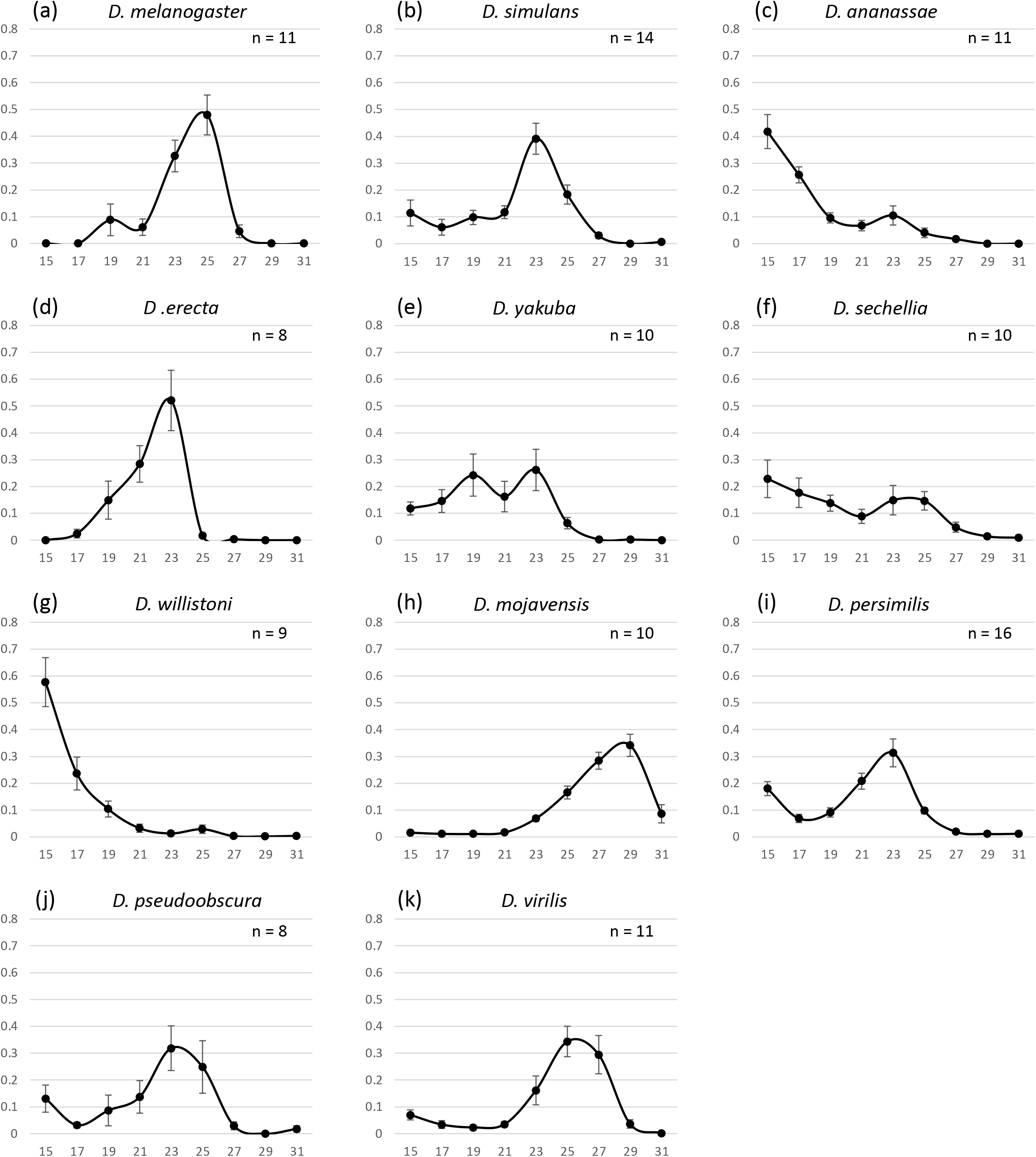
Temperature preference of 11 *Drosophila* species. (a) ∼ (k) represents the distribution of flies on the temperature gradient field in each species; (a) *D. melanogaster*, (b) *D. simulans*, (c) *D. ananassae*, (d) *D. erecta*, (e) *D. yakuba*, (f) *D. sechellia*, (g) *D. willistoni*, (h) *D. mojavensis*, (i) *D. persimilis*, (j) *D. pseudoobscura*, and (k) *D. virilis*. Percentages (y axis) of the flies positioned in nine temperature ranges (x axis) are shown (see Methods). Error bars indicate standard error of means.

Then, we determined the preferred temperature (see Methods) for each species (Fig. 6) and compared it with the optimal temperature required for locomotor activity (the temperature at which the species showed the highest activity) (Fig. 1, Supplementary Fig.2 and Fig. 6). The comparison revealed that the preferred temperatures almost coincided with the optimal temperature required for locomotor activity in *D. melanogaster, D. mojavensis, D. persimilis*, and *D. pseudoobscura*. However, the preferred temperature was higher than the optimal temperature in *D. simulans* and *D. virilis* and lower than the optimal temperature in *D. ananassae, D. yakuba, D. willistoni*, and *D. sechellia*. These results indicated that *Drosophila* species do not always prefer the temperature at which their locomotor activity levels are high.

**Figure 6.**
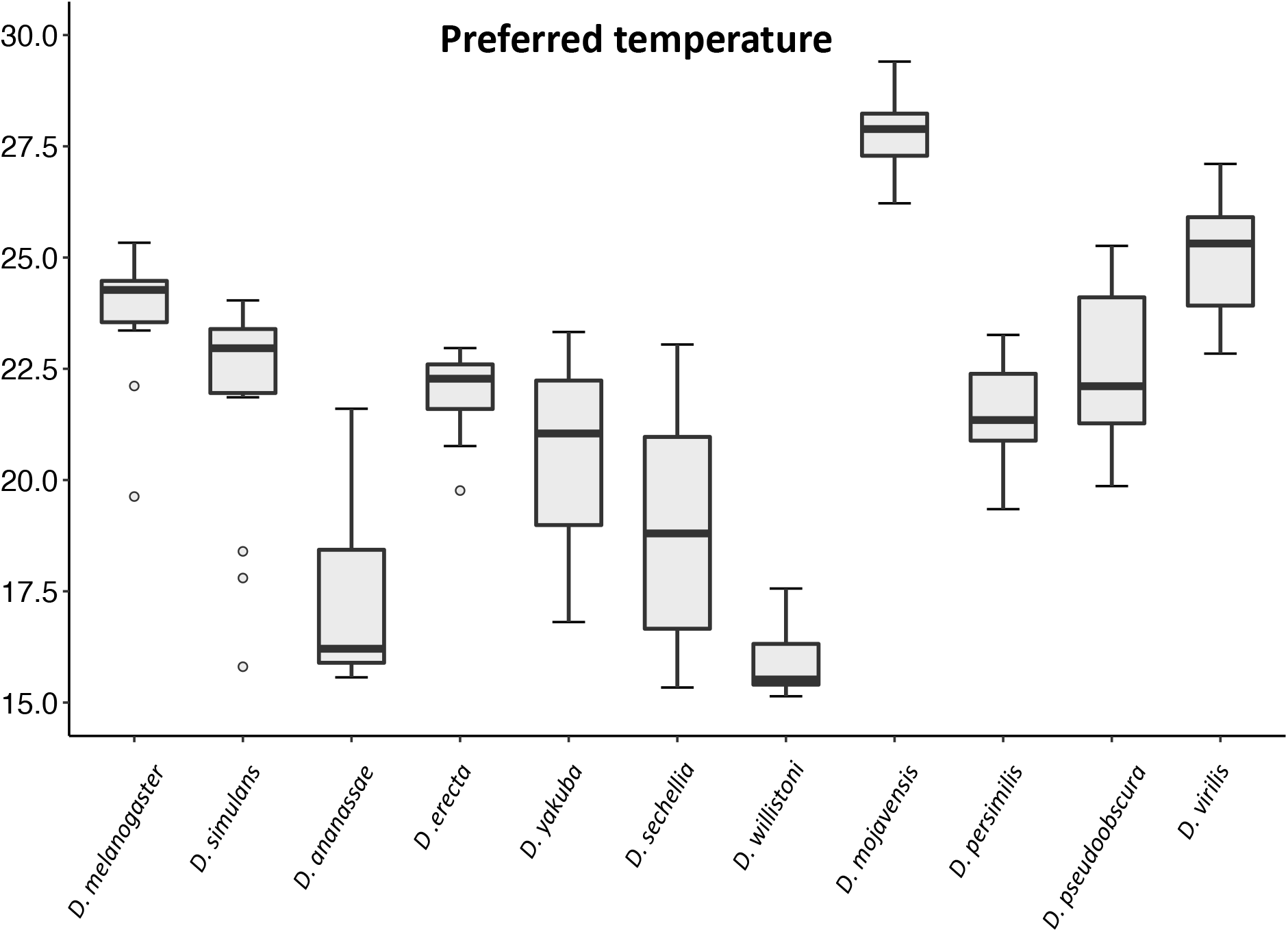
Preferred temperatures of 11 *Drosophila* species. Preferred temperatures of 11 *Drosophila* species are shown. Median of the preferred temperature of each species is as follows; *D. melanogaster*; 24.3°C, *D. simulans*; 23.0°C, *D. ananassae*; 16.2°C, *D. erecta*; 22.3°C, *D. yakuba*; 21.0°C, *D. sechellia*; 18.8°C, *D. willistoni*; 16.0°C, *D. mojavensis*; 27.9°C, *D. persimilis*; 21.3°C, *D. pseudoobscura*; 21.1°C, and *D. virilis*; 25.3°C. Circles with box plots show outliers.

### Function of antennae in terms of temperature preference behavior

In *D. melanogaster*, studies have experimentally demonstrated that antennae sense the external cold temperature and are indispensable for cold avoidance and temperature preference behavior [23, 29]. Therefore, we investigated the function of antennae in the temperature preference behavior of the 11 species using antenna ablation experiments (Fig. 7). Consistent with previous studies, *D. melanogaster* with antenna ablation was broadly distributed to cooler temperatures at ≤ 23°C. However, intriguingly, antenna ablation had no such effects in the other species. In *D. erecta* and *D. virilis*, the distribution peaks became smoother without large shifts of peak temperatures. In contrast, the peaks became prominent in *D. willistoni* and *D. persimilis*. However, heat avoidance at higher temperature (31°C) appeared to be lost in *D. mojavensis*. The other species showed no drastic changes by antenna ablation. These results suggest that the function of antennae in the temperature preference behavior can be diverse among *Drosophila* species.

**Figure 7.**
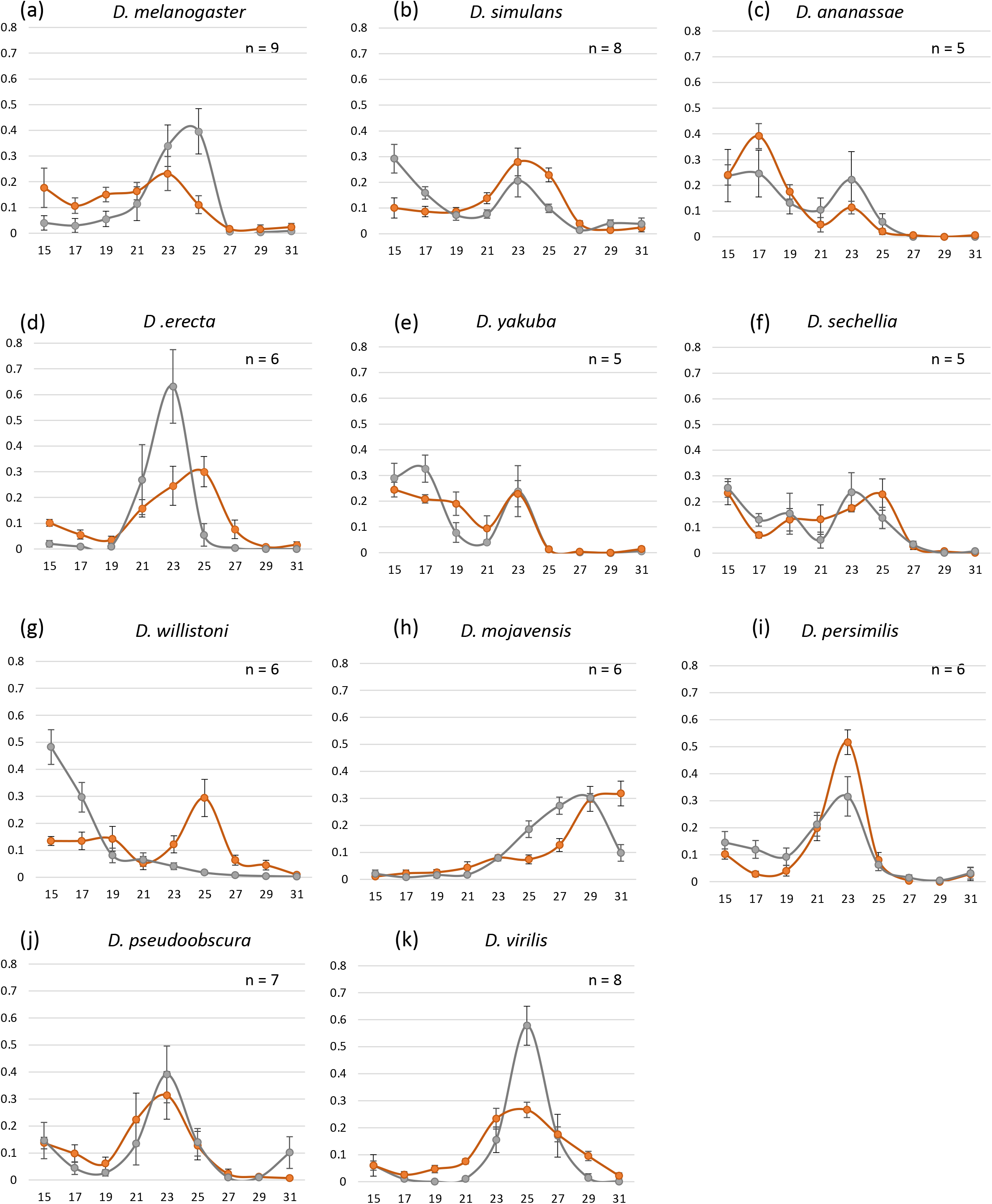
Temperature preference of 11 *Drosophila* species with antenna ablation. (a) ∼ (k) represents the distribution of control (gray line) and antenna ablated (orange line) flies on the temperature gradient field in each species; (a) *D. melanogaster*, (b) *D. simulans*, (c) *D. ananassae*, (d) *D. erecta*, (e) *D. yakuba*, (f) *D. sechellia*, (g) *D. willistoni*, (h) *D. mojavensis*, (i) *D. persimilis*, (j) *D. pseudoobscura*, and (k) *D. virilis*. Error bars indicate standard error of means.

To compare the effect of the antenna quantitatively, we determined the preferred temperature in flies with antenna ablation (Fig. 8). We found that ablation had no significant effect on the preferred temperature in eight species (Mann–Whitney U test: *p* < 0.05). In only three species, namely, *D. melanogaster, D. simulans*, and *D. willistoni*, ablation had a significant impact on the preferred temperature (Mann–Whitney U test: *p* < 0.05). The preferred temperature shifted to a cooler temperature by ablation in *D. melanogaster* and *D. simulans*. In contrast, the preferred temperature shifted to a warmer temperature in *D. willistoni*. The shifts of the preferred temperature to a cooler or warmer side by antenna ablation suggest the loss of cold avoidance (or heat attraction) and heat avoidance (or cold attraction), respectively. Collectively, these results suggest that the antenna-dependent regulation of the temperature preference behavior observed in *D. melanogaster* is not the typical system.

**Figure 8.**
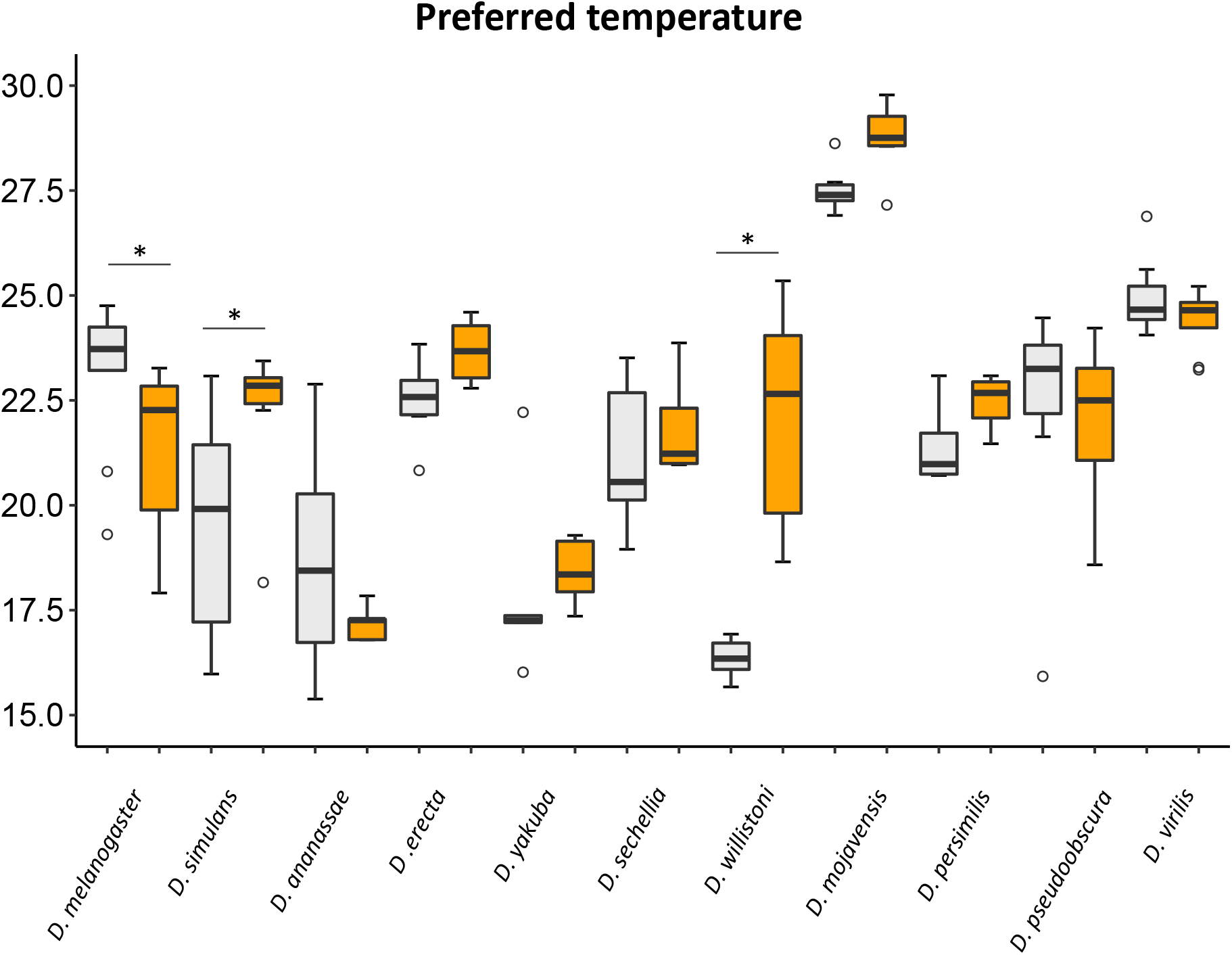
Effects of the antenna ablation on preferred temperatures. Preferred temperatures in control (gray box) and antenna ablated (orange box) flies are shown. Horizonal bars with * above the box indicate significant difference between control and antenna ablation flies (Mann–Whitney U tests: *p* < 0.05).

## Discussion

We investigated the effects of temperature on locomotor activities and temperature preference behavior in 11 *Drosophila* species. As per our findings, we observed that these species exhibited species-specific activity levels and sleep duration. Most species exhibited higher locomotor activities at temperatures they encountered in their habitats. Temperature had a unique impact on the amount of total daily activities and sleep in a species-specific manner. All species exhibited high activities around morning and/or evening, irrespective of temperature. Some species changed the amount of daily activity maintaining the daytime/nighttime activity ratio, whereas others changed the total activity changing the daytime/nighttime activity ratio. With regard to temperature preference behavior, all species avoided higher temperatures; however, the degree of avoidance or attraction to cold temperature was diverse among species, resulting in species-specific responses to the temperature gradient. Furthermore, we experimentally found that the function of the antenna in the temperature preference behavior was not common among *Drosophila* species, and some species identified the preferred temperature without antennae. These results suggest that these *Drosophila* species have acquired species-specific temperature response systems based on their own life strategy to cope with ambient temperatures.

Previous studies have showed that some characteristics of life history of *D. melanogaster* are adapted in laboratory conditions by laboratory culture [30, 31]. We used the laboratory strains for all analyses in this study. Therefore, it is uncertain how much the observed species-specific characteristics in locomotor activity and temperature-preference behavior can explain the species-specific adaptation to the natural thermal environment. On the other hand, it has also been reported that laboratory maintenance did not eclipse fundamental species-specific ecological and physiological characteristics in *Drosophila* species [32]. It is reasonable to assume that some characteristics are altered while others are maintained by laboratory maintenance. To understand the extent of species-specific characteristics maintenance in laboratory strains and the presence of intraspecific variation in certain *Drosophila* species, we need to comparatively analyze many different strains, including freshly-established different local strains and other laboratory-maintained strains. Previously, the laboratory strains of 11 *Drosophila* species have been used for comparative studies of the embryonic development and the larval locomotor behavior [33, 34]. Interestingly, the correlation between species-specific characteristics and their natural habitats has been reported with the laboratory strains [34]. Therefore, the experimental results using laboratory strains, including our study, should be valuable benchmarks for future comparative analyses in understanding the exact species-specific characteristics and adaptations.

Increases in locomotor activity would be related to opportunities for mating and foraging. Therefore, the temperature at which the species is most active is considered to be optimal for activity in the species. In this study, excluding *D. melanogaster*, all the other species exhibited significant or remarkable differences in total activity among the analyzed temperatures, and the temperature at which high total activity was observed matched the temperature in the habitat. This result suggests that these species are adapted to the temperature zone of their habitats. On the other hand, *D. melanogaster* could maintain the same amount of daily activity at various temperatures. A previous study showed that effect of temperature on the locomotor activity was different between *D. melanogaster* strains established from different climate zones; activity levels at temperature 17ºC to 29ºC were about the same for the strain established from the tropical climate zone, though the activity was decreased at 17ºC compared with 25ºC and 29ºC in the temperate strain [13]. Additionally, it has also been reported that the spontaneous activity was higher at 24ºC than at 17ºC in another temperate strain [35]. Furthermore, it has also been shown that the temperature of the developmental period significantly affects the activity level in an adult [36]. Therefore, it is noteworthy whether such plastic features are specific to *D. melanogaster*, and whether these are related to the fact that this species is cosmopolitan.

Regarding the daily locomotor activity pattern in these species, we were able to note a species-specific effect of temperature on the daily activity pattern. It is well known that *D. melanogaster* has two daily activity surges, known as morning and evening peaks [37]. As in *D. melanogaster*, we found that each species has a morning and evening peak for activity at the optimal temperature with the species-specific pattern. Some species were more active in the evening, whereas some species were more active in the morning. These differences might correlate with the niche in time, which is generally observed in animals [38]. Interestingly, the shape of each peak and the ratio of surge magnitude are species-specific, and they retain their characteristics even at different temperatures in most cases. Moreover, each species has its unique range of total daily activity, which is diverse among species (Supplementary Fig. 1). Because the species-specific daily activity pattern and level might be tightly correlated with the species-specific lifestyle and life strategy, they would be robustly maintained even at different temperatures.

In addition to locomotor activity, we observed that sleep was affected by temperature. In most species, total daily sleep length was lesser at the temperature at which total daily activity was higher. In this study, the definition of sleep for *D. melanogaster* was applied to all other species, i.e., an immobile state for ≥5 minutes [39]. However, the total locomotor activity was significantly different among species. Several species, such as *D. erecta* and *D. mojavensis*, were much less active compared with *D. melanogaster*. Therefore, it should be considered whether applying the definition for *D. melanogaster* to all other species is suitable.

If the optimal temperature for activity is the most adequate temperature for certain species, the preferred temperature should match the optimal temperature for activity. As anticipated, the four species, namely, *D. melanogaster, D. mojavensis, D. persimilis*, and *D. pseudoobscura*, preferred the optimal activity temperature or nearby temperatures in the temperature preference experiments. However, there were discrepancies between the preferred and optimal temperatures for the remaining species. Four tropical or subtropical species, i.e., *D. ananassae, D. yakuba, D. sechellia*, and *D. willistoni*, preferred temperatures lower than the optimal activity temperature, whereas a cosmopolitan species, *D. simulans*, and a temperate species, *D. virilis*, preferred temperatures higher than the optimal activity temperature. It can be assumed that there are several factors affecting the temperature preference behavior. Temperature can influence various behaviors and physiological phenomena such as courtship, egg-laying, energy storage, and energy efficiency [40-43]. Additionally, it has been reported that the rearing temperature and ages affect the adult temperature preference in several *Drosophila* species [44, 45]. If the optimal temperature is different among these traits, the temperature that flies prefer might differ depending on situations they are placed and the behavior or phenomenon that is precedent. Moreover, their temperature preference may be affected by noxious stress [17]. Because tropical and subtropical species have increased chances of being exposed to higher temperature in their habitats, they might have a higher adaptive plasticity for heat avoidance and/or cold attraction. For the same reason, some temperate species might be adapted to avoiding cold stress and/or heat attraction. The preference behavior of a species would be determined under the balance of these factors, and also the priority factor would vary among species.

Previous studies have shown that antennal thermosensing regulates cold avoidance in *D. melanogaster* [19, 22, 23]. Therefore, it is expected that the temperature preference behavior of *Drosophila* species is also largely dependent on antennal thermosensing. However, in this study, we found that antenna ablation affected temperature sensing in only *D. simulans* and *D. willistoni*, in addition to *D. melanogaster*. The effects of ablation were diverse among these species. For example, in *D. willistoni*, antenna-ablated flies showed cold avoidance unlike *D. melanogaster*, suggesting that antennal thermosensing functions to attract flies to cold. The temperature preference behavior is a type of decision-making behavior made by specific brain neurons to stay at or leave from the current temperature regime [46]. In *D. melanogaster*, manipulating the subsets of dopaminergic neurons has altered the temperature preference behavior [47]. Hence, it is possible that the difference in the effect of antenna ablation between these two species is caused by the difference in the manner of utilizing temperature information from the antenna for the decision-making behavior rather than the difference in the properties of antennae as a temperature sensor.

Unexpectedly, we found no significant contribution of antennae for the temperature preference behavior in 6 of the 11 examined species. This result indicates that these species do not use antennae for regulating the temperature preference behavior but utilize other temperature sensors. Previous studies have shown that the anterior cell (AC) neurons inside the head sense innocuous warmth above 25°C and exert essential function in the temperature preference behavior in *D. melanogaster* [19]. However, the multiple dendritic (md) neurons in the adult body wall are candidates to sense noxious warmth for the escape behavior in adults, although it has not yet been tested [18]. Therefore, in addition to adult AC cells, md neurons are can be considered potential candidates to sense temperature for the temperature preference behavior in these species. As insects commonly have thermosensors in antennae with specific morphologies [48], it is difficult to believe that the antennae of these species do not sense temperature. In these species, the decision-making for temperature preference would not be largely dependent on thermal information from the antenna thermosensors. *Drosophila* species would make a species-specific decision using multiple sources of thermal information depending on the type of behaviors, resulting in their unique life strategy for adaptation to the temperature environment in their habitats.

In this investigation, by conducting comparative studies on 11 *Drosophila* species, we found that the effects of temperature on behavior are diverse and that there is an interspecific variation in the responses to temperature. This finding suggests that even at the same environmental temperature, different species respond differently to temperature, and it is favorable for some species but unfavorable for others. This phenomenon would be associated with the difference in behavioral output following the sensing of temperature information. Moreover, these differences may be associated with strategies for adaptation to habitat temperature conditions. Understanding the mechanisms responsible for these differences would shed light on the adaptation strategy and evolution of species. This study was able to elucidate the diversity of temperature adaptation in *Drosophila* species and opened avenues for understanding the underlying mechanisms behind their diverse temperature adaptations.

## Methods

### Fly strains

The following fly strains were used in this study: *D. melanogaster* (Canton-S, k-aba006), *D. simulans* (Rakujuen, k-abb292), *D. ananassae* (AABBg1, k-s01), *D. erecta* (k-s02), *D. yakuba* (k-s03), *D. sechellia* (k-s10), *D. willistoni* (k-s13), *D. mojavensis* (k-s15), *D. persimilis* (k-s11), *D. pseudoobscura* (k-s12), and *D. virilis* (k-s14) from KYORIN-Fly, *Drosophila* Species Stock Center in Kyorin University. All strains were reared on a standard cornmeal medium in a 12-h/12-h light/dark (LD) cycle. The rearing temperature was 20°C for *D. persimilis*, but it was 23°C for the others.

### Measurements of locomotor activity and sleep

To measure daily locomotor activities, male adult flies were collected within 4 h after eclosion. Before applying the activity tubes, the collected flies were maintained in standard medium for 1–2 days. Under ice anesthetization, an individual fly was applied into monitoring tubes (7mm diameter for *D. virilis* and 5mm diameter for others) filled with 5% sucrose (Wako) and 2% Bacto agar (BD) on one side, and activity was evaluated using the *Drosophila* Activity Monitor (DAM) system (TriKinetics Inc.) under the 12-h/12-h LD cycle condition (light on 8 AM and off 8 PM every day) at five different temperatures (17°C, 20°C, 23°C, 26°C, and 29°C). After 2 days of acclimation for each recording temperature, the locomotor activities were recorded every 1 min for 3 days. The average total daily locomotor activity of an individual fly was calculated using the total counts for 3 days for each recording. Each sample size is shown in Supplementary Fig. 3. The median value of total daily locomotor activity in each species was calculated with data on total daily activities at all experimental temperatures (Supplementary Fig. 1). The daily activity pattern was estimated by summarizing activity counts every 30 min (Fig. 2 and Supplementary Fig. 3). The total activities of first and second half of 24 h and daytime and nighttime were analyzed by summarizing the counts for 12 h from 2 AM to 2 PM and 2 PM to 2 AM (Supplementary Fig. 10) and from 8 AM to 8 PM and 8 PM to 8 AM (Fig. 3), respectively.

The definition of sleep in *D. melanogaster*, i.e., 5 or more minutes of immobile state [26], was applied for this analysis (Fig. 4 and Supplementary Fig. 7). We determined sleep behavior based on the activity data recorded using the DAM system. The average total daily sleep duration was calculated using the total sleep duration for 3 days for each recording. The number of sleep events per day and the average length of single sleep were also calculated using the same datasets.

### Temperature preference assay and antenna ablation experiment

To analyze the temperature preference behavior, we designed the temperature gradient field according to a previous study with some modifications [49]. In this assay, the air temperature between a plexiglass cover and an aluminum plate was monitored using a FLUKE 52II thermometer with multiple temperature probes (FLUKE 80PK-1) at 4 or 6 points along the temperature gradient field. Before applying the flies, the plexiglass cover was coated with super Rain-X (Rain-X) to prevent the flies from climbing up the cover. The tests were conducted in a room at approximately 23°C and a relative humidity of approximately 70%–85% under light illumination. Because *D. mojavensis* randomly gathered around the corners of the plexiglass cover under light illumination, the tests were conducted under red light condition for this species. The temperature gradients were set between 14°C and 32°C. For the tests, 30–50 flies (flies aged 3–7 days reared with 12-h/12-h LD cycles) were applied into this apparatus without anesthetizing, and after 60 min, the distribution of flies was recorded by taking photos. The temperatures of all flies were determined after recording; the positions of all flies in the field were determined using the ImageJ point tool, and the temperature at which each fly was positioned was determined using two adjacent temperature probes. The measurements were performed between 14:00 and 20:00 and repeated at least 8 times using different individuals.

To analyze the distribution of flies on the temperature gradient field, the flies were classified into nine temperature ranges based on a method described in a previous study [19, 21]. Flies with <16°C and >30°C were categorized as the 15°C and 31°C temperature range of flies, respectively. Flies with 16°C–30°C were categorized into seven different temperature ranges with an interval of 2°C. The percentage of flies positioned in each temperature range was analyzed in each assay. The mean percentage of distribution and the standard error in each temperature range were calculated (Figs. 5 and 7). Based on the individual fly temperature data, the median temperature was calculated in each assay as the preferred temperature (Figs. 6 and 8).

For antenna ablation experiments, the second and/or third antenna segments were ablated from newly eclosed flies (within 4 h after eclosion) using pointed tweezers under ice anesthesia. In this experiment, ice-anesthetized flies were used for experimental controls (Figs. 7 and 8). After operation or anesthetization (control), flies were maintained for 4–7 days before conducting the temperature preference behavior test. The measurements were performed between 14:00 and 20:00 and repeated at least 5 times using different individuals.

### Statistical analysis

All statistical analyses were performed in R, version 3.6.1. Statistical analyses of the effects of temperatures on daily activities and temperature preference were conducted using nonparametric Mann–Whitney U tests for pairwise comparisons (Figs. 3 and 8 and Supplementary Figs. 4 and 10) and Kruskal–Wallis tests with Bonferroni/Dunn test [50] for multiple pair comparisons (Fig. 2 and Supplementary Figs. 1, 4, 8–10, 12–13). Correlation analyses among behaviors were performed using Spearman’s rank correlation coefficient (Supplementary Table 1). In order to estimate the optimal temperatures and maximum performance of locomotor activity, we fitted 3 or 13 TPC models using *rTPC* package for R [51] and close the best fitted one based on the lowest Akaike Information Criteria (AIC) values (Supplementary Fig. 2).

## Supporting information

All supplemental information

## Data availability

The datasets generated during and/or analyzed during the current study are available from the corresponding author on reasonable request.

## Acknowledgements

We greatly thank to laboratory members for technical assistance, Dr. Masahito T. Kimura for comments and critical reading of the manuscript, Dr. Taro Ueno for introduction of DAM system, Dr. Daniel Padfield for comments about *rTPC* analyses, KYORIN-Fly for fly lines, Enago for editing the manuscript. This work was supported by grants from the Ministry of Education, Culture, Sports, Science and Technology of Japan (KAKENHI grants 16H04658 to T.A.).

## Author contributions

Conceptualization, F. I. and T. A.; Methodology, F. I. and T. A.; Formal Analysis, F. I.; Investigation, F. I. and T. A.; Writing-Original Draft & Editing, F. I. and T. A.; Funding Acquisition, T.A.; Supervision, T. A.

## References

1. Franks, S. J. & Hoffmann, A. A. Genetics of climate change adaptation. Annu. Rev. Genet. 46, 185–208 (2012).

2. Kimura, M. T. Adaptations to temperate climates and evolution of overwintering strategies in the Drosophila melanogaster species group. Evolution 42, 1288–1297 (1988).

3. Overgaard, J. & MacMillan, H. A. The integrative physiology of insect chill tolerance. Annu. Rev. Physiol. 79, 187–208 (2017).

4. Gibert, P., Moreteau, B., Pétavy, G., Karan, D. & David, J. R. Chill-coma tolerance, a major climatic adaptation among Drosophila species. Evolution 55, 1063–1068 (2001).

5. Toda M. J. (2014) DrosWLD-Species: Taxonomic information database for world species of Drosophilidae [updated 29 May 2022]. URL: https://bioinfo.museum.hokudai.ac.jp/db/modules/stdb/.

6. O’Grady, P. M. & DeSalle, R. Phylogeny of the genus Drosophila. Genetics 209, 1–25 (2018).

7. Andersen, J. L. et al. How to assess Drosophila cold tolerance. Funct. Ecol. 29, 55–65 (2015).

8. Kellermann, V. et al. Upper thermal limits of Drosophila are linked to species distributions and strongly constrained phylogenetically. Proc. Nat. Acad. Sci. USA 109, 16228–16233 (2012).

9. Kimura, M. T. Quantitative response to photoperiod during reproductive diapause in the Drosophila auraria species-complex. J. Insect Phys. 36, 147 (1990).

10. Markow, T. A. & O’Grady, P. Reproductive ecology of Drosophila. Funct. Ecol. 22, 747–759 (2008).

11. Xue, Q., Majeed, M. Z., Zhang, W. & Ma, C. Adaptation of Drosophila species to climate change — A literature review since 2003. J. Integr. Agric. 18, 805–814 (2019).

12. Watanabe, K. et al. Interspecies comparative analyses reveal distinct carbohydrate-responsive systems among Drosophila Species. Cell Rep. 28, 2594-2607.e7 (2019).

13. Klepsatel, P. & Gáliková, M. Developmental temperature affects thermal dependence of locomotor activity in Drosophila. J. Therm. Biol. 103, 103153 (2022).

14. Dubowy, C. & Sehgal, A. Circadian rhythms and sleep in Drosophila melanogaster. Genetics 205, 1373–1397 (2017).

15. Low, K. H., Lim, C., Ko, H. W. & Edery, I. Natural variation in the splice site strength of a clock gene and species-specific thermal adaptation. Neuron 60, 1054–1067 (2008).

16. Brown, E. B. et al. Variation in sleep and metabolic function is associated with latitude and average temperature in Drosophila melanogaster. Ecol. Evol. 8, 4084 (2018).

17. Dillon, M. E., Wang, G., Garrity, P. A. & Huey, R. B. Review: Thermal preference in Drosophila. J. Therm. Biol. 34, 109–119 (2009).

18. Barbagallo, B. & Garrity, P. A. Temperature sensation in Drosophila. Curr. Opin. Neurobiol. 34, 8–13 (2015).

19. Hamada, F. N. et al. An internal thermal sensor controlling temperature preference in Drosophila. Nature 454, 217–220 (2008).

20. Hong, S. et al. Histamine and its receptors modulate temperature-preference behaviors in Drosophila. J. Neurosci. 26, 7245–7256 (2006).

21. Sayeed, O. & Benzer, S. Behavioral genetics of thermosensation and hygrosensation in Drosophila. Proc. Nat. Acad. Sci. USA 93, 6079–6084 (1996).

22. Ni, L. et al. A gustatory receptor paralogue controls rapid warmth avoidance in Drosophila. Nature 500, 580–584 (2013).

23. Budelli, G. et al. Ionotropic receptors specify the morphogenesis of phasic sensors controlling rapid thermal preference in Drosophila. Neuron 101, 738-747.e3 (2019).

24. Kellis, M. et al. Evolution of genes and genomes on the Drosophila phylogeny. Nature 450, 203–218 (2007).

25. Chiu, J. C., Low, K. H., Pike, D. H., Yildirim, E. & Edery, I. Assaying locomotor activity to study circadian rhythms and sleep parameters in Drosophila. J. Vis. Exp. 43, e2157. doi:10.3791/2157 (2010).

26. Hall, J. C. Genetics and molecular biology of rhythms in Drosophila and other insects. Adv. Genet. 48, 1–280 (2003).

27. Buhr, E., Yoo, S. & Takahashi, J. S. Temperature as a universal resetting cue for mammalian circadian oscillators. Science 30, 379–385 (2010).

28. Parisky, K., Agosto Rivera, J., Donelson, N., Kotecha, S. & Griffith, L. Reorganization of sleep by temperature in Drosophila requires light, the homeostat, and the circadian clock. Curr. Biol. 26, 882–892 (2016).

29. Gallio, M., Ofstad, T. A., Macpherson, L. J., Wang, J. W. & Zuker, C. S. The coding of temperature in the Drosophila Brain. Cell 144, 614–624 (2011).

30. Hoffmann, A. A., Hallas, R., Sinclair, C. & Partridge, L. Rapid loss of stress resistance in Drosophila melanogaster under adaptation to laboratory culture. Evolution 55, 436–438 (2001).

31. Sgrò, C. M. & Partridge, L. Laboratory adaptation of life history in Drosophila. Am. Nat. 158, 657–658 (2001).

32. Maclean, H. J., Kristensen, T. N., Sørensen, J. G. & Overgaard, J. Laboratory maintenance does not alter ecological and physiological patterns among species: a Drosophila case study. J. Evol. Biol. 31, 530–542 (2018).

33. Kuntz, S. G. & Eisen, M. B. Drosophila embryogenesis scales uniformly across temperature in developmentally diverse species. PLoS Genet. 10, e1004293 (2014).

34. Matsuo, Y., Nose, A. & Kohsaka, H. Interspecies variation of larval locomotion kinematics in the genus Drosophila and its relation to habitat temperature. BMC Biol. 19, 176–4 (2021).

35. Plantamp, C. et al. Phenotypic plasticity in the invasive pest Drosophila suzukii: activity rhythms and gene expression in response to temperature. J. Exp. Biol. 222, jeb199398. doi: 10.1242/jeb.199398 (2019).

36. MacLean, H. J., Kristensen, T. N., Overgaard, J., Sørensen, J. G. & Bahrndorff, S. Acclimation responses to short-term temperature treatments during early life stages causes long lasting changes in spontaneous activity of adult Drosophila melanogaster. Physiol. Entomol. 42, 404–411 (2017).

37. Grima, B., Chelot, E., Xia, R. & Rouyer, F. Morning and evening peaks of activity rely on different clock neurons of the Drosophila brain. Nature 431, 869–873.

38. Taylor, R. S. & Friesen, V. L. The role of allochrony in speciation. Mol. Ecol. 26, 3330–3342 (2017).

39. Shaw, P. J., Cirelli, C., Greenspan, R. J. & Tononi, G. Correlates of sleep and waking in Drosophila melanogaster. Science 287, 1834–1837 (2000).

40. Berrigan, D. & Partridge, L. Influence of temperature and activity on the metabolic rate of adult Drosophila melanogaster. Comparative biochemistry and physiology. Comp. Biochem. Physiol. A Physiol. 118, 1301–1307(1997).

41. Klepsatel, P., Wildridge, D. & Gáliková, M. Temperature induces changes in Drosophila energy stores. Sci. Rep. 9, 5239 (2019).

42. Schnebel, E. M. & Grossfield, J. Mating-temperature range in Drosophila. Evolution 38, 1296–1307 (1984).

43. Smith, J. M. The Effects of temperature and of egg-laying on the longevity of Drosophila subobscura. J. Exp. Biol. 35, 832 (1958).

44. MacLean, H. J. et al. Temperature preference across life stages and acclimation temperatures investigated in four species of Drosophila. J. Therm. Biol. 86, 102428 (2019).

45. Rajpurohit, S. & Schmidt, P. S. Measuring thermal behavior in smaller insects: A case study in Drosophila melanogaster demonstrates effects of sex, geographic origin, and rearing temperature on adult behavior. Fly 10, 149–161 (2016).

46. Bang, S. et al. Dopamine signalling in mushroom bodies regulates temperature-preference behaviour in Drosophila. PLoS Genet. 7, e1001346 (2011).

47. Buhl, E., Kottler, B., Hodge, J. J. L. & Hirth, F. Thermoresponsive motor behavior is mediated by ring neuron circuits in the central complex of Drosophila. Sci. Rep. 11, 155–9 (2021).

48. Foelix, R. F., Stocker, R. F. & Steinbrecht, R. A. Fine structure of a sensory organ in the arista of Drosophila melanogaster and some other dipterans. Cell Tissue Res. 258, 277–287 (1989).

49. Goda, T., Leslie, J. R. & Hamada, F. N. Design and analysis of temperature preference behavior and its circadian rhythm in Drosophila. J. Vis. Exp. 83, e51097. doi: 10.3791/51097 (2014).

50. Dinno A. dunn.test: Dunn’s test of multiple comparisons using rank sums. (2017). URL: http://cran.r-project.org/package=dunn.test.

51. Padfield, D., O’Sullivan, H. & Pawar, S. rTPC and nls.multstart: A new pipeline to fit thermal performance curves in R. tMethods Ecol. Evol. 12 1138–1143 (2021).

