## Supplementary material for "Comparative analysis of temperature preference behavior and effects of temperature on daily behavior in eleven *Drosophila* species": All supplemental information

### Supplementary Information

Supplemental Figure 1.

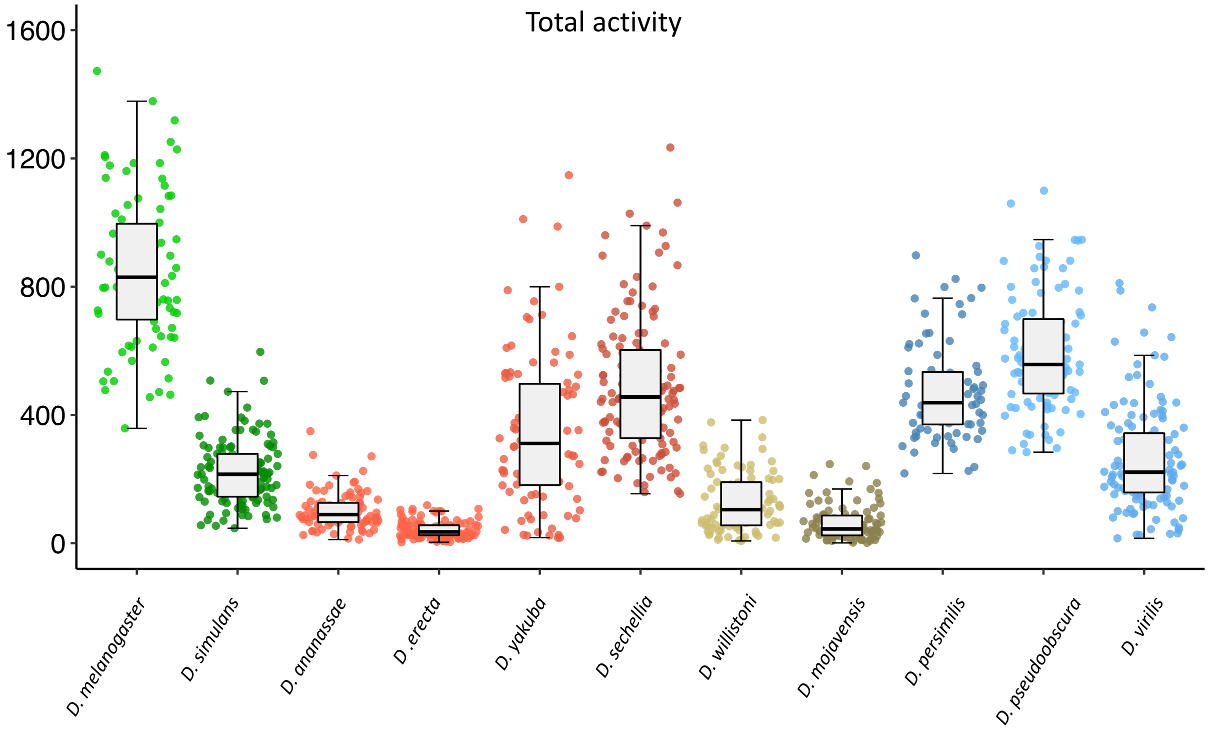

Supplemental Figure 2.

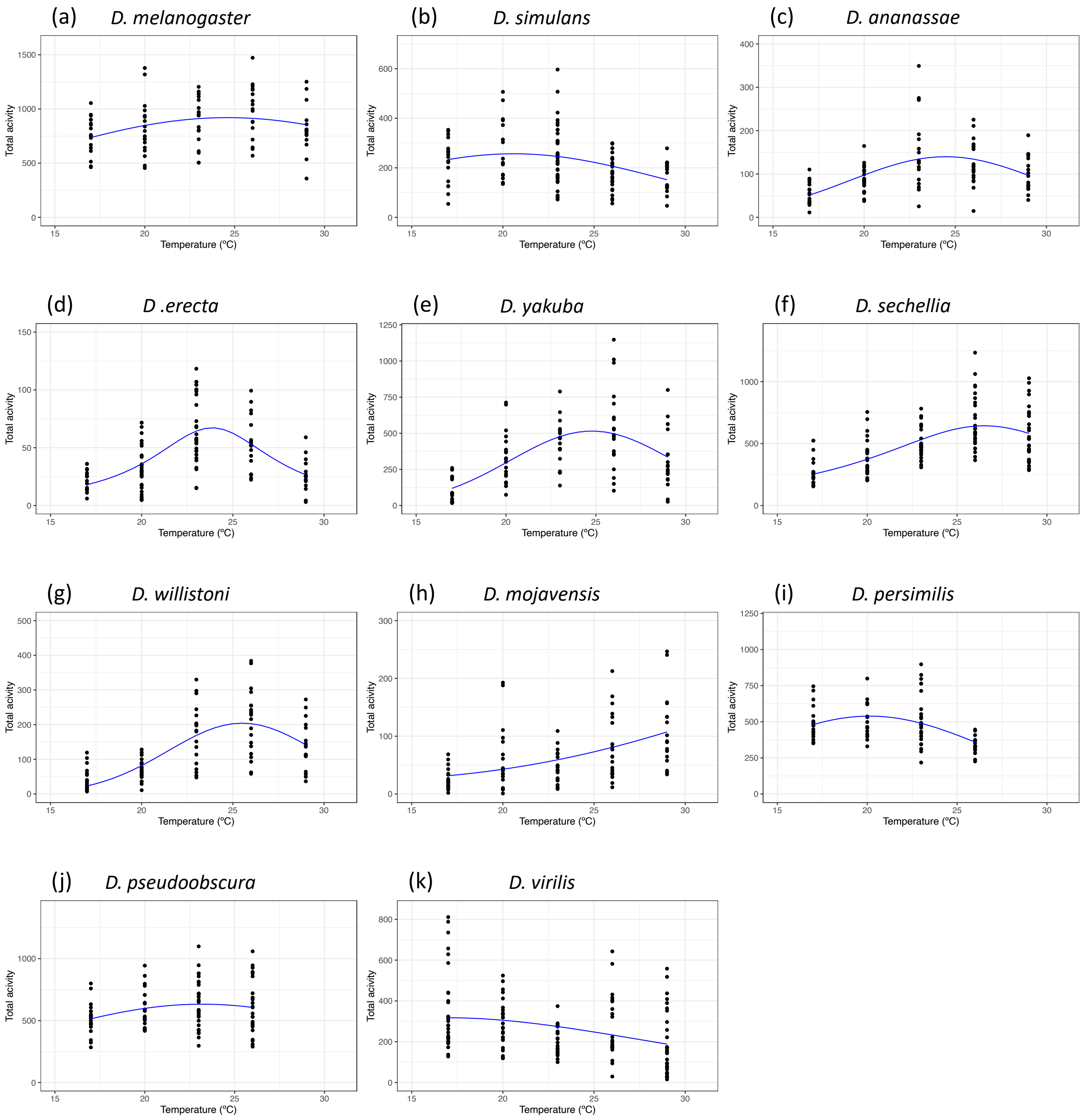

Supplementary Figure 3 (a)

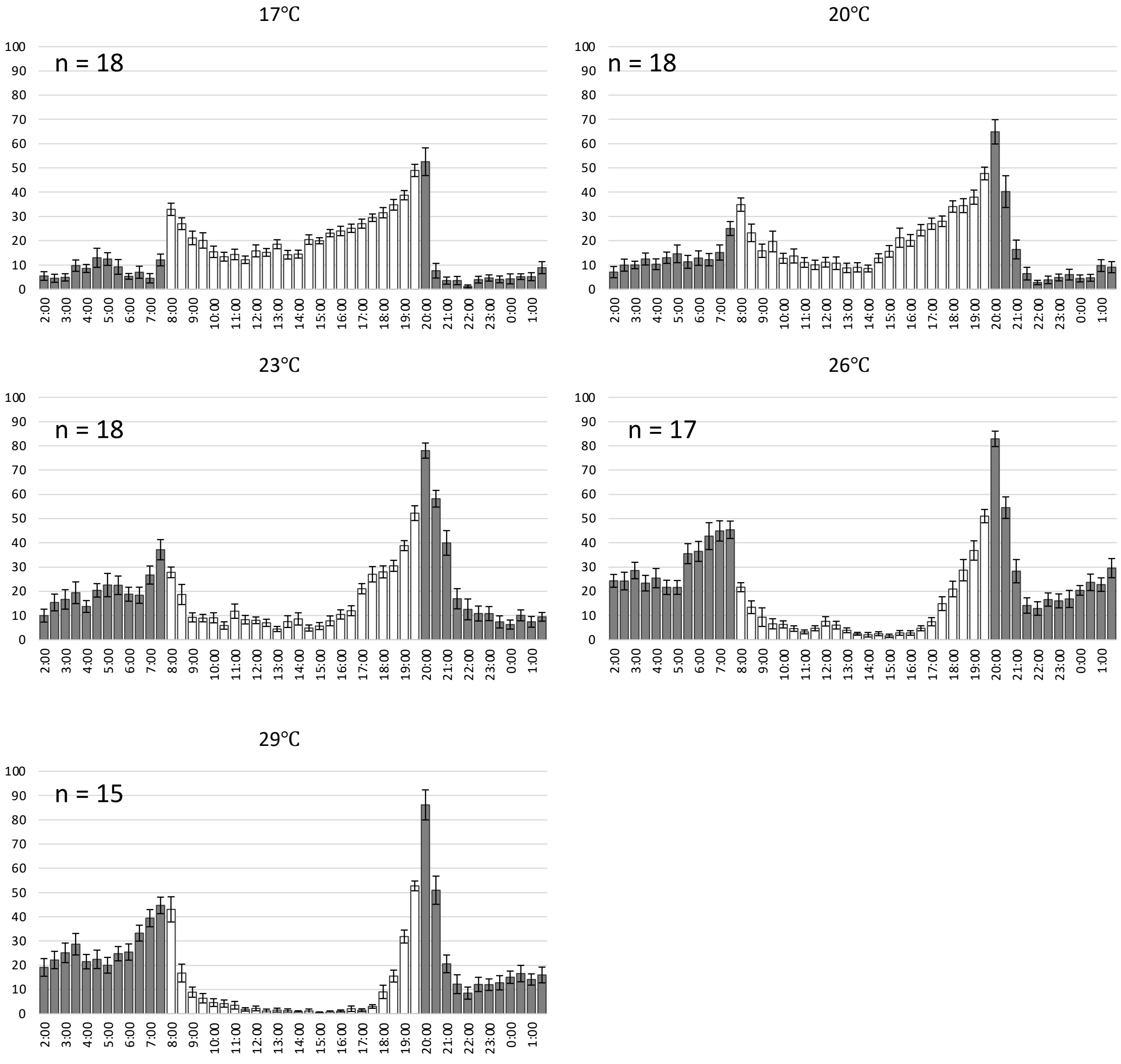


*D. melanogaster*

Supplementary Figure 3 (b)

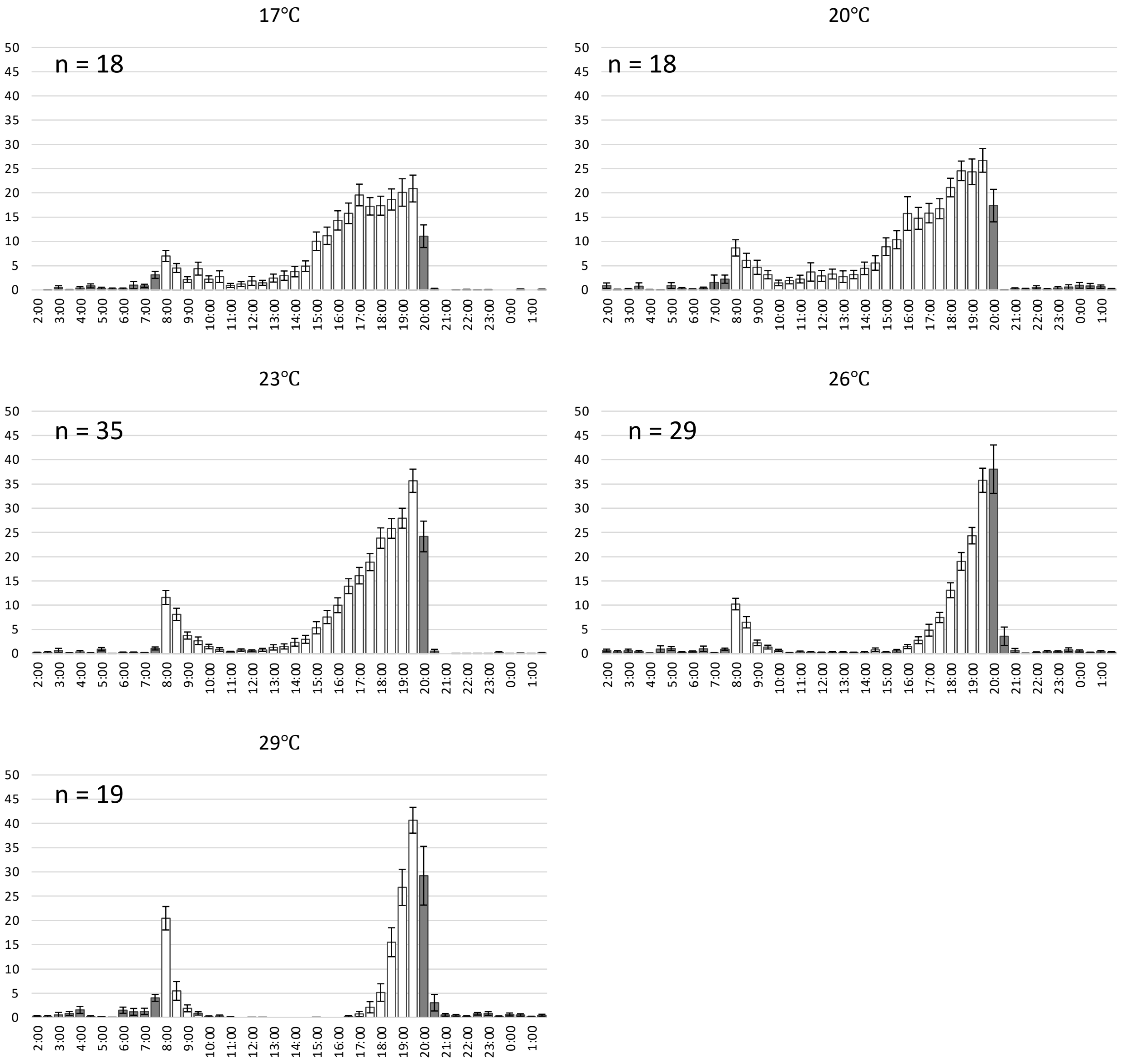


*D. simulans*

Supplementary Figure 3 (c)

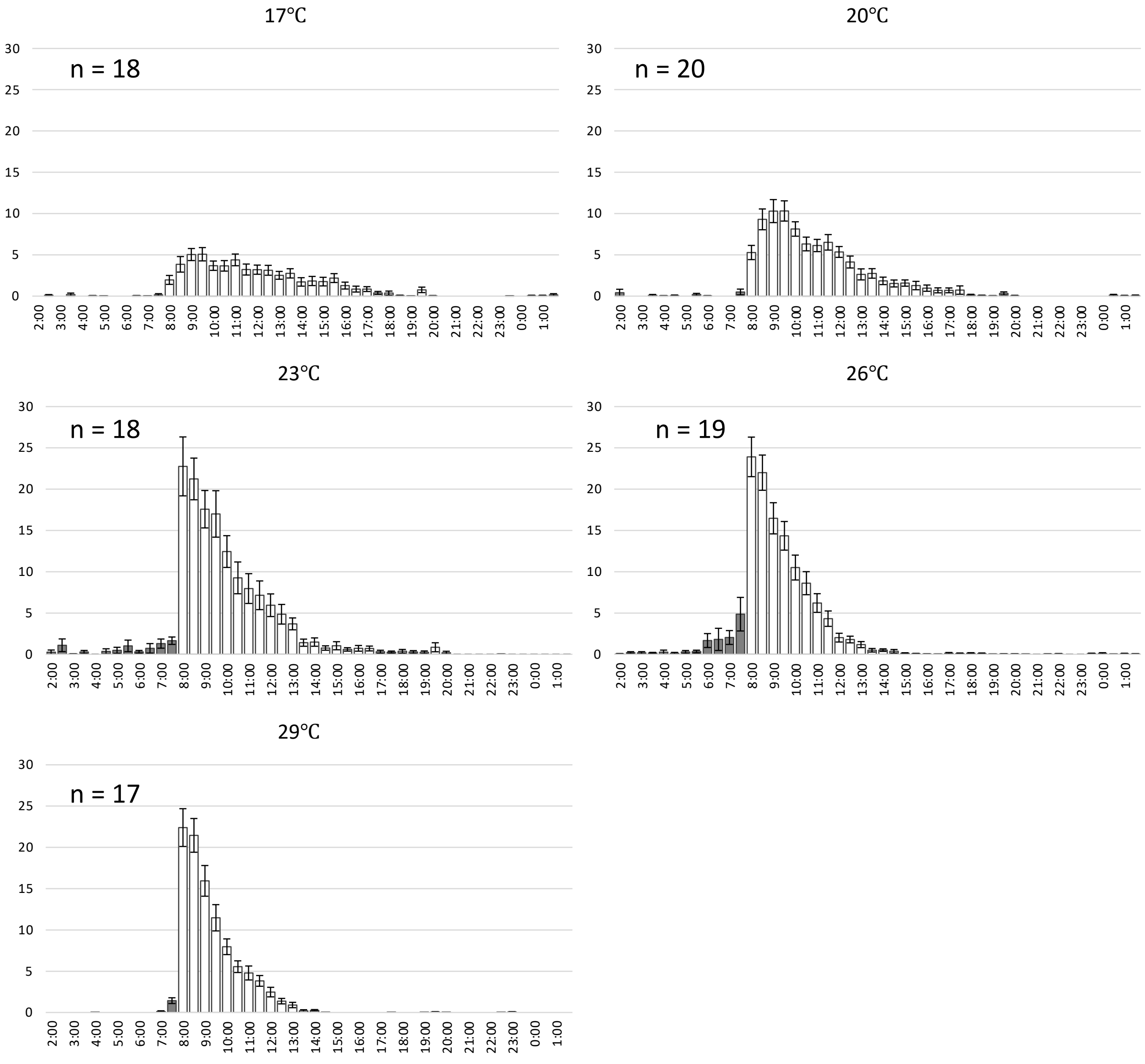


*D. ananassae*

Supplementary Figure 3 (d)

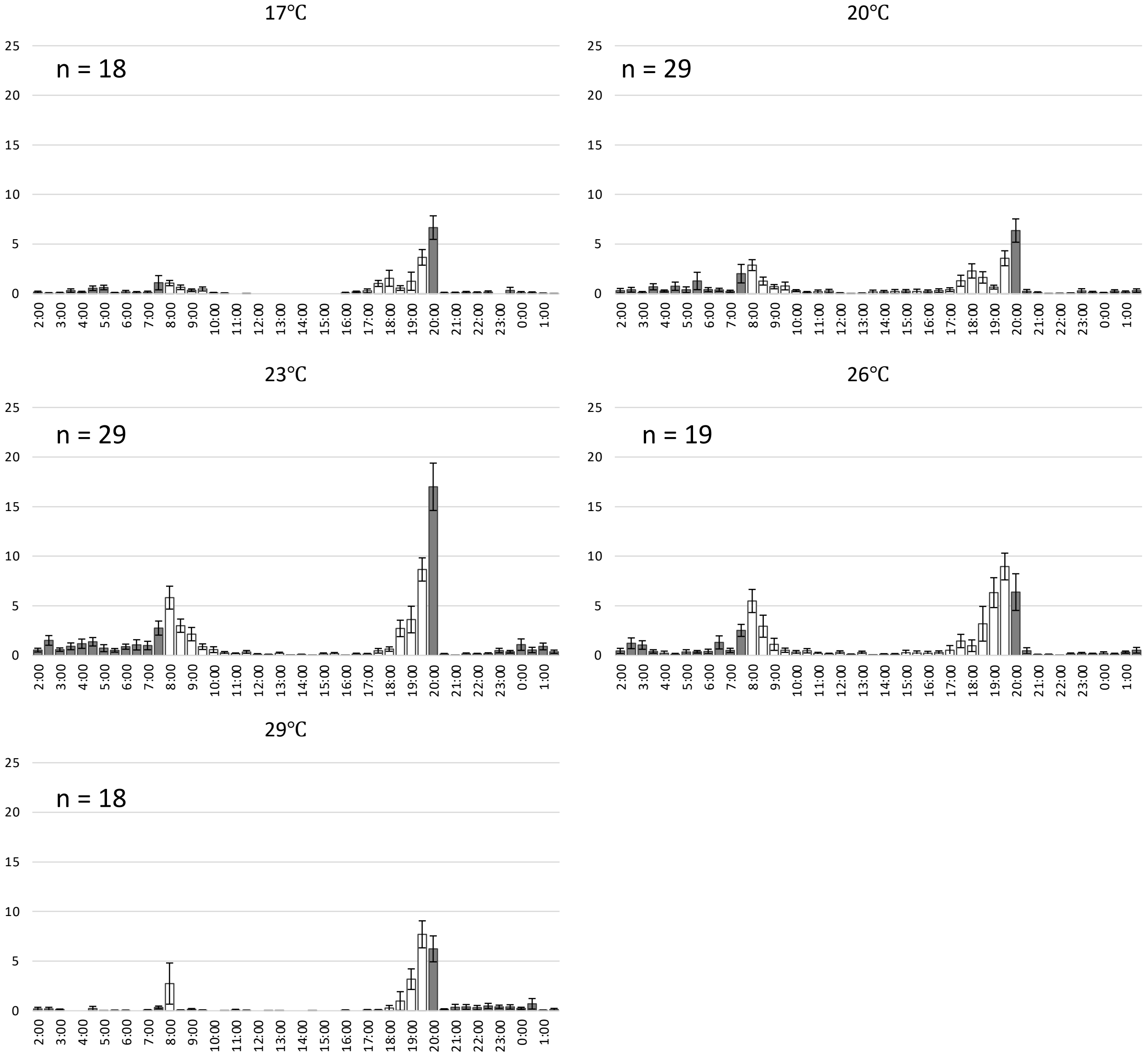


*D. erecta*

#### Supplementary Figure 3 (e)

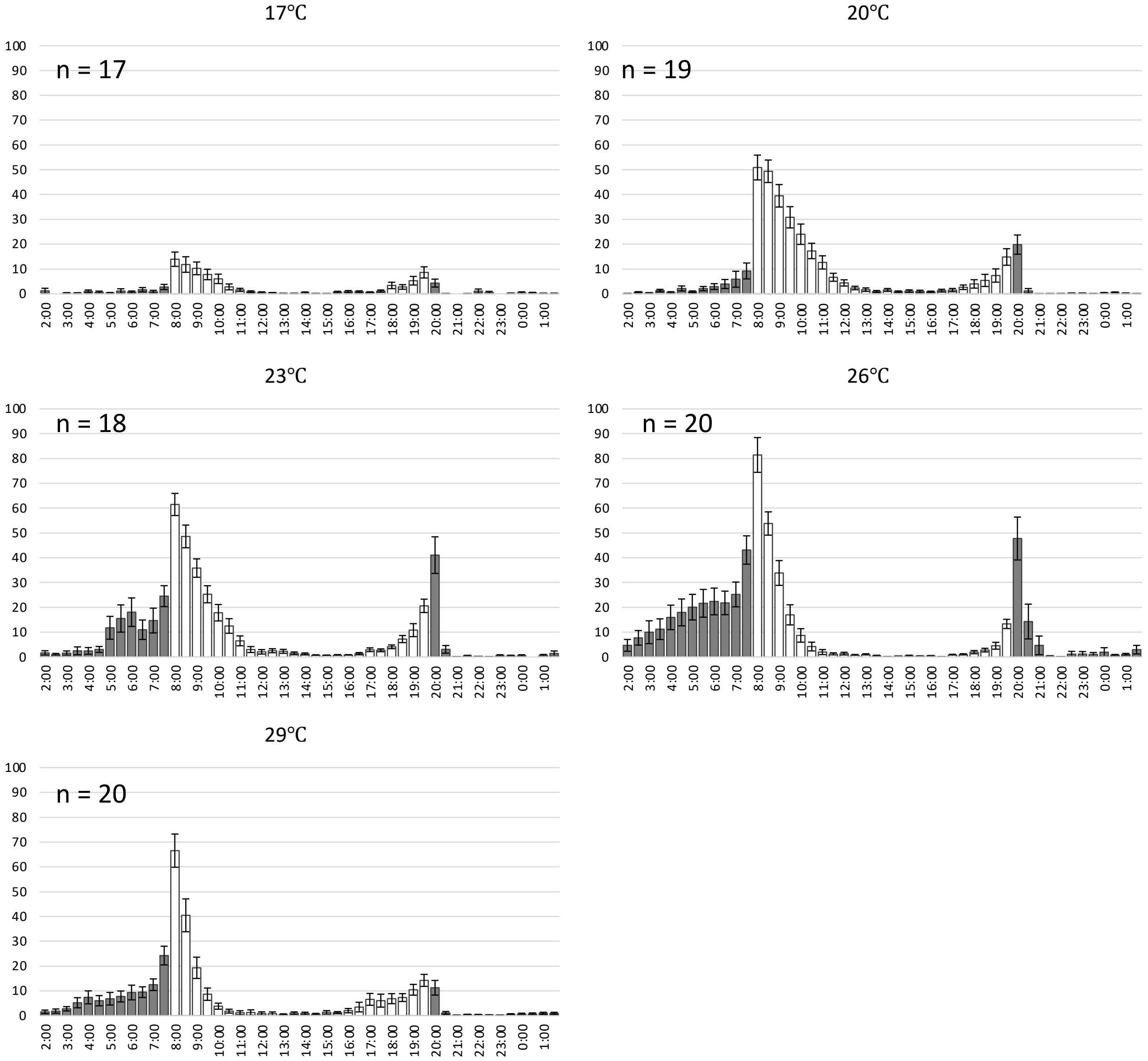


*D. yakuba*

#### Supplementary Figure 3 (f)

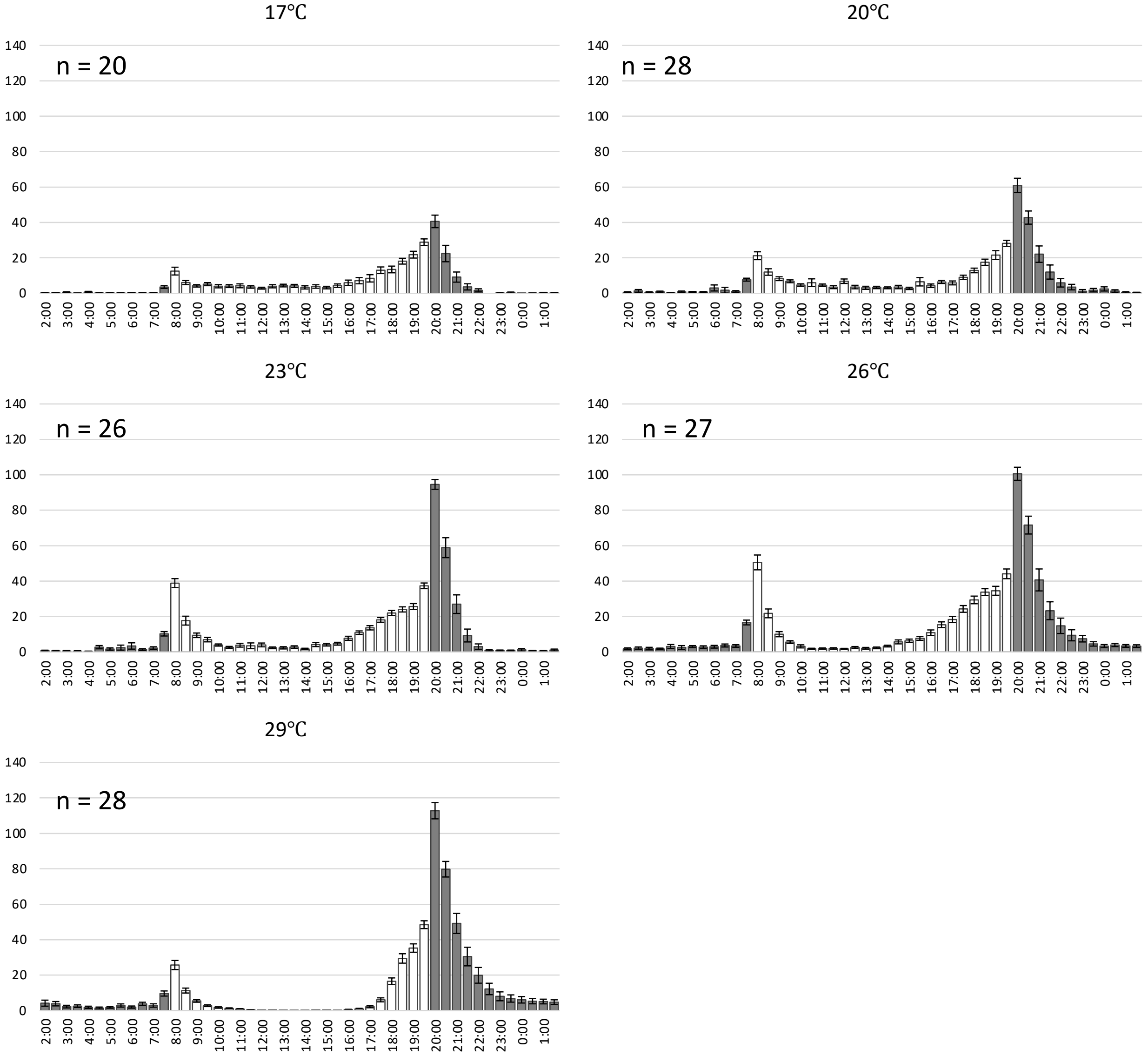


*D. sechellia*

Supplementary Figure 3 (g)

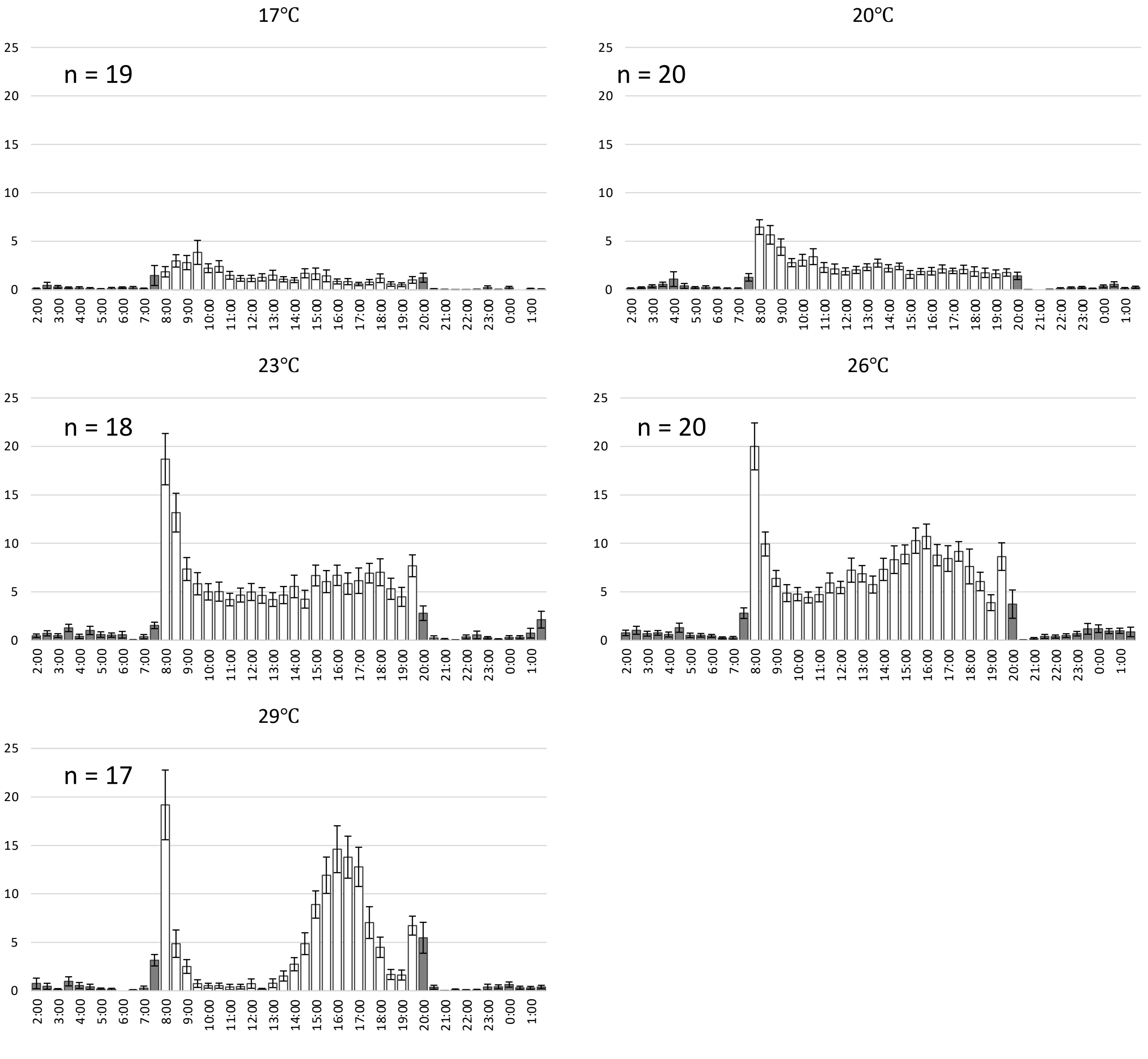


*D. willistoni*

Supplementary Figure 3 (h)

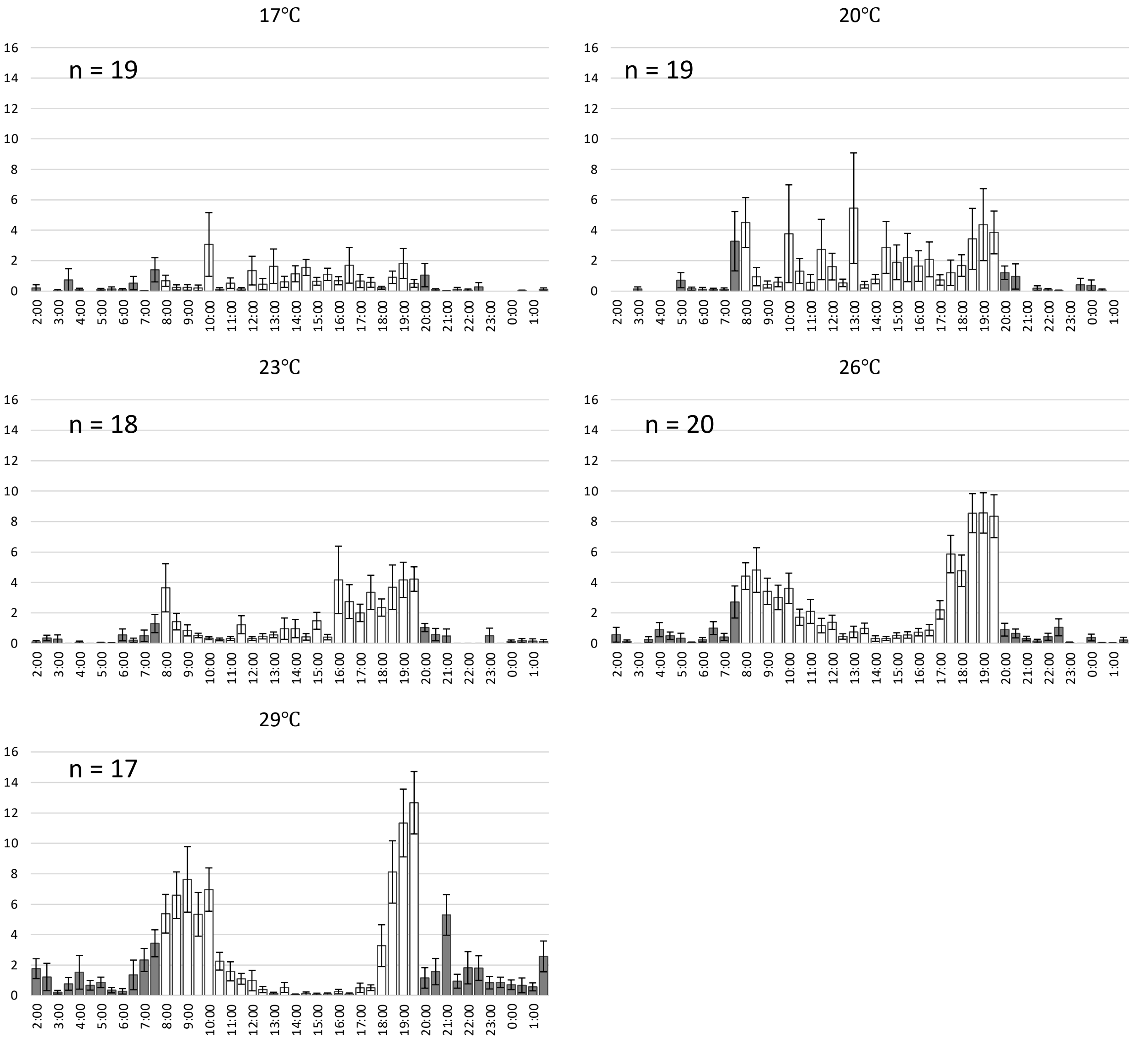


*D. mojavensis*

Supplementary Figure 3 (i)

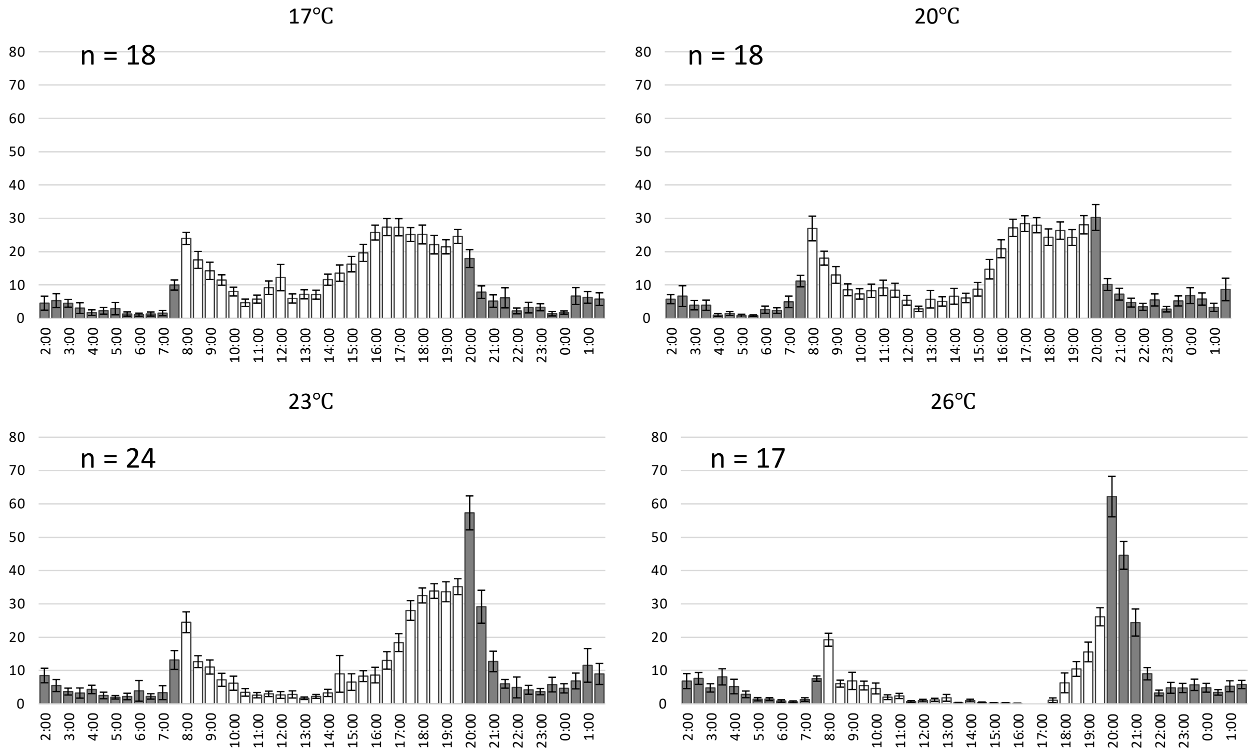


*D. persimilis*

Supplementary Figure 3 (j)

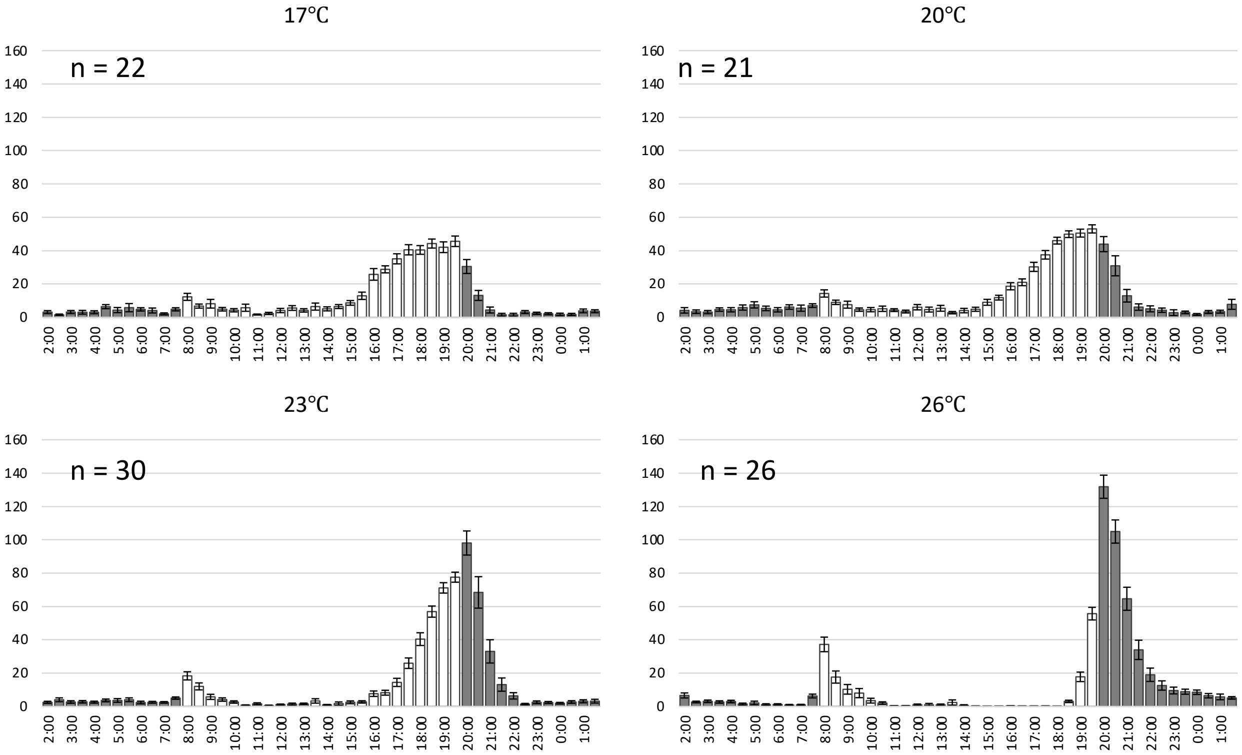


*D. pseudoobscura*

Supplementary Figure 3. (k)

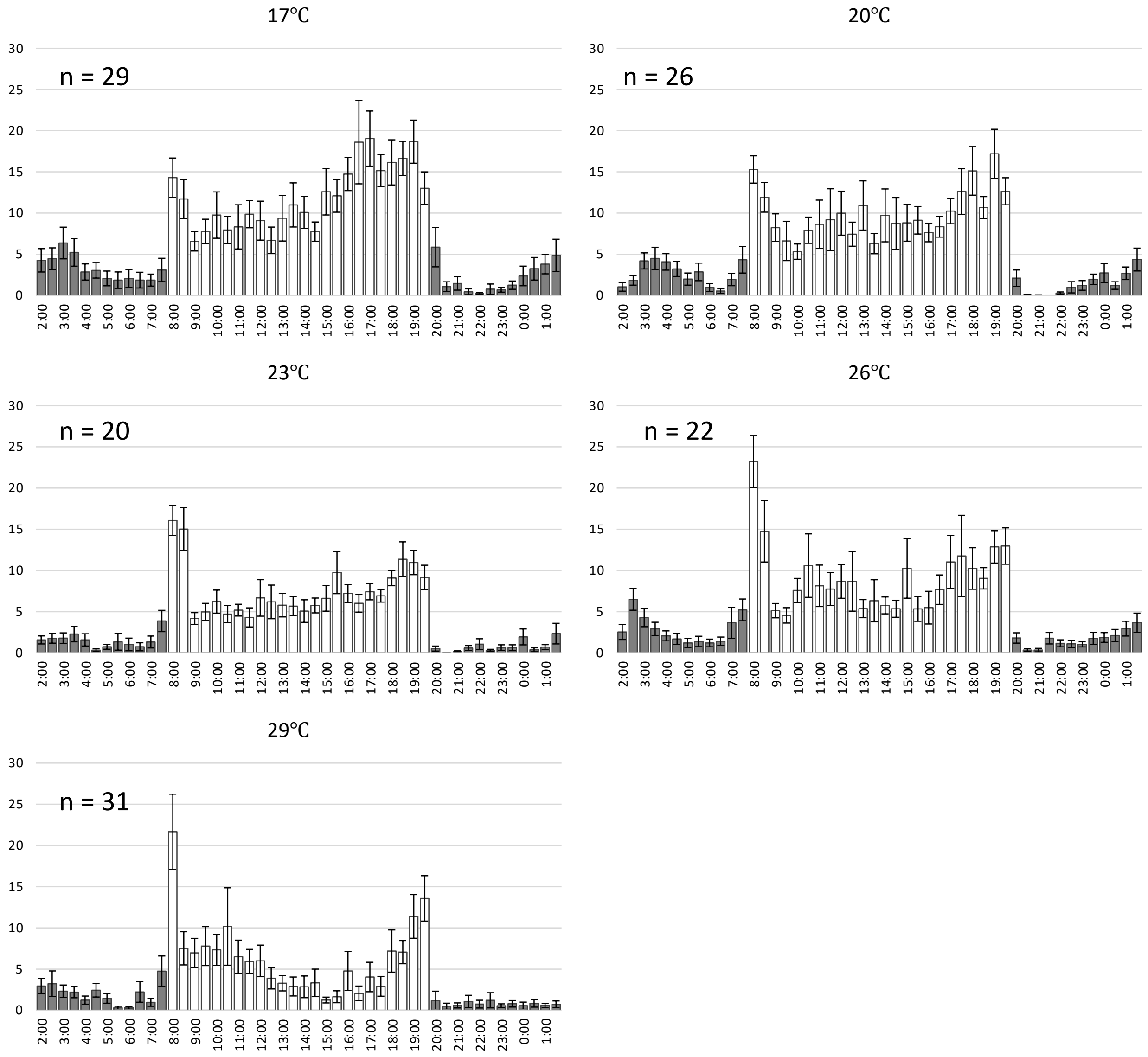


*D. virilis*

Supplementary Figure 4.

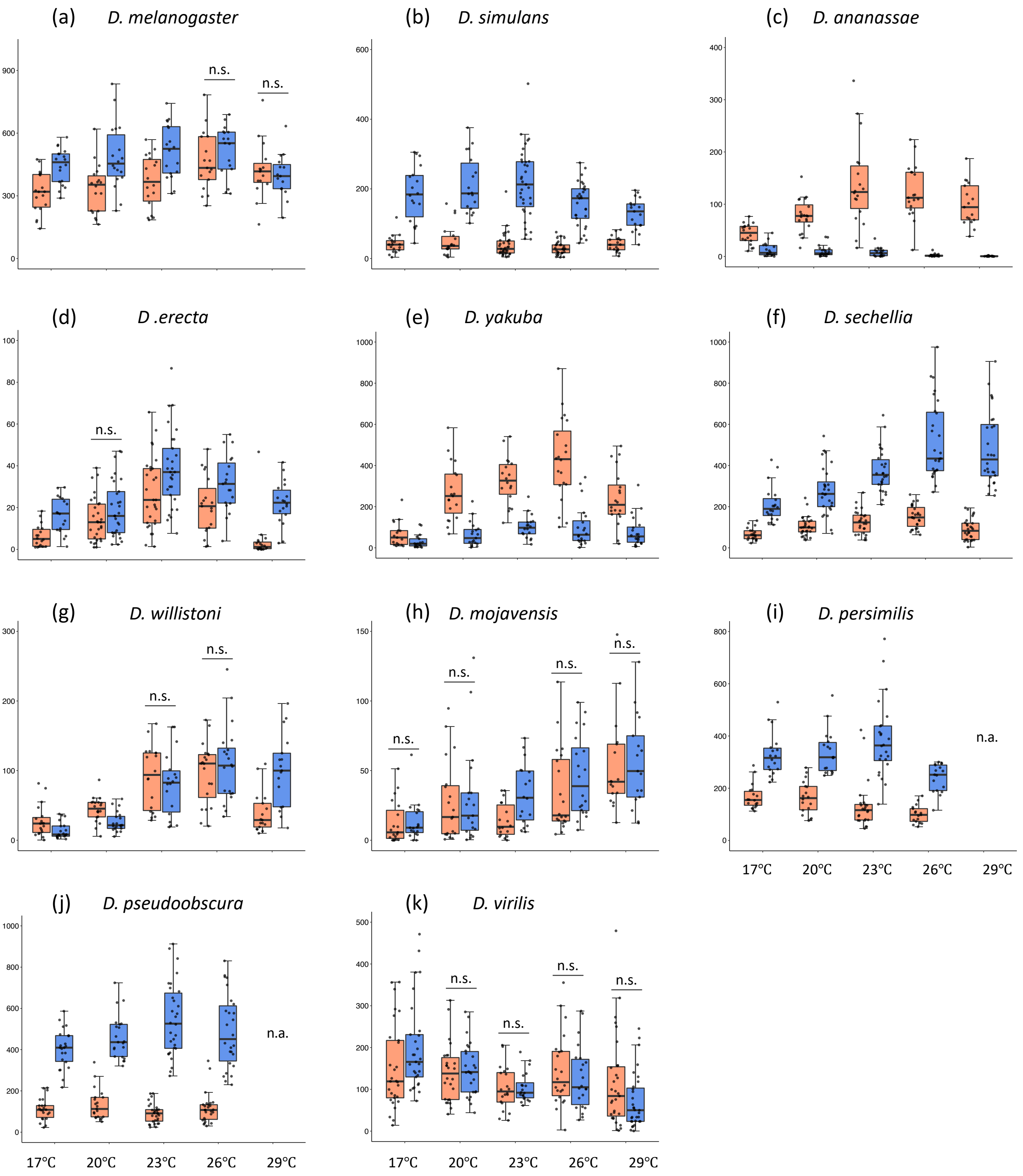

Supplementary Figure 5.

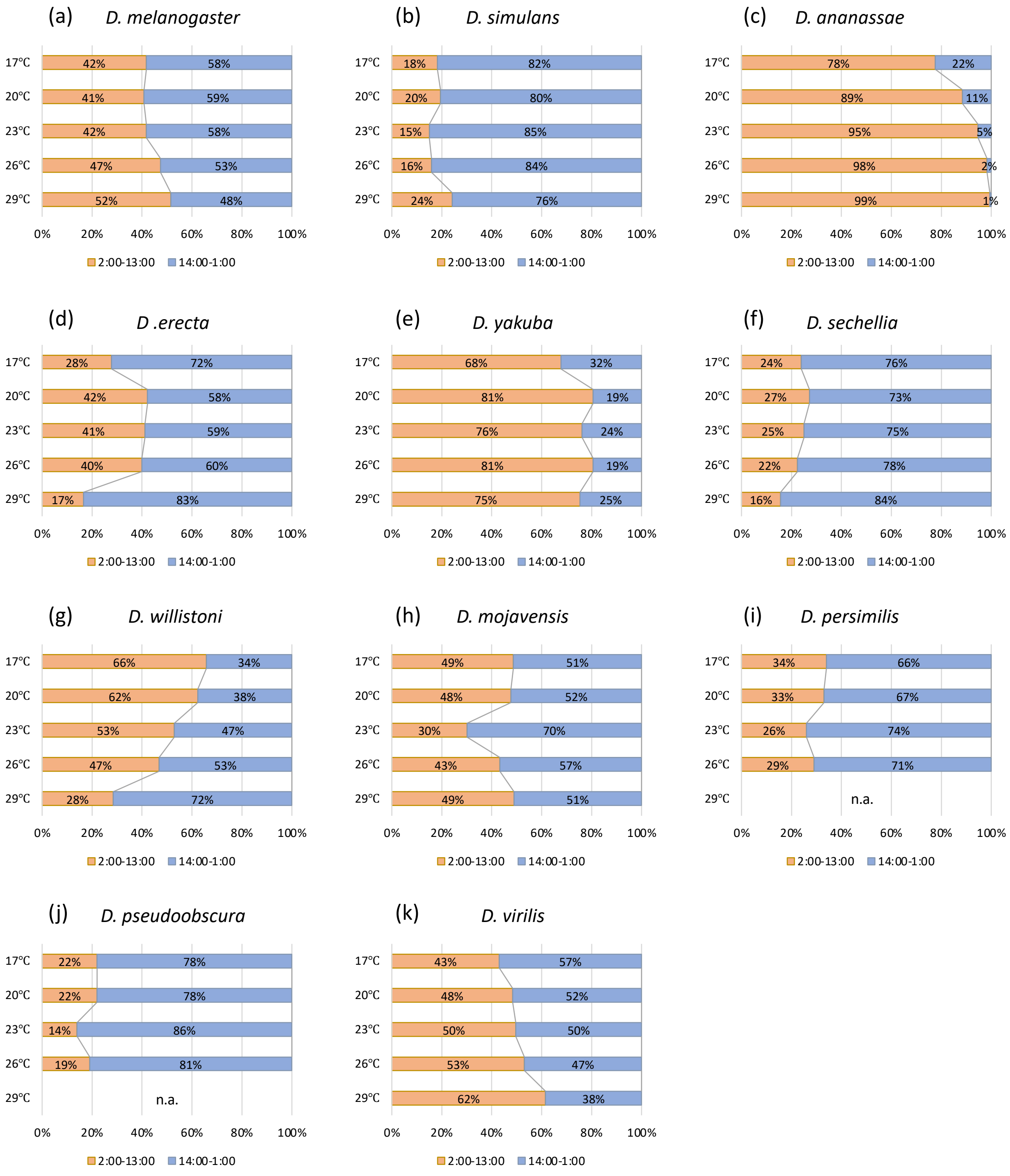

#### Supplementary Figure 6.

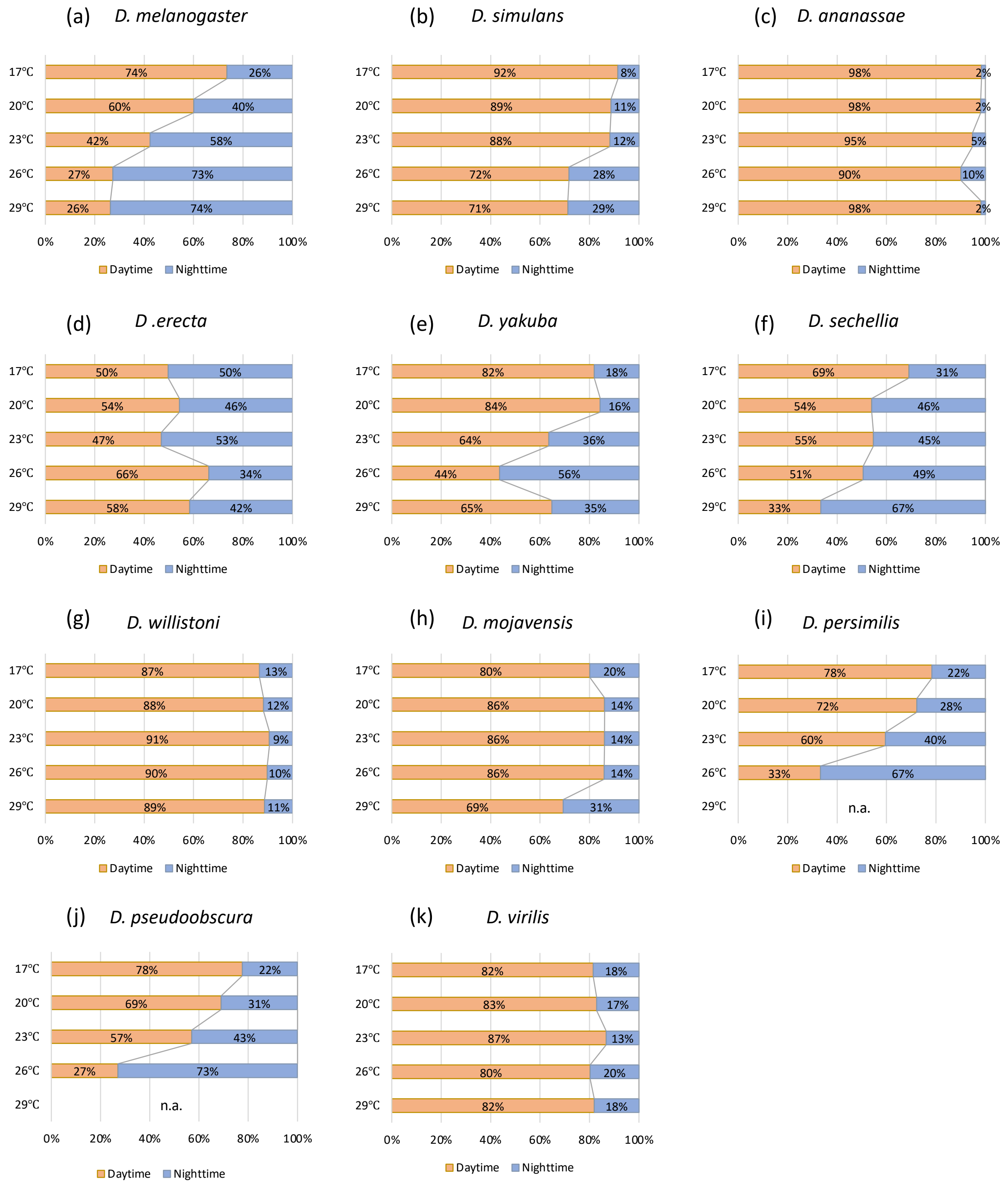

Supplementary Figure 7. (a)

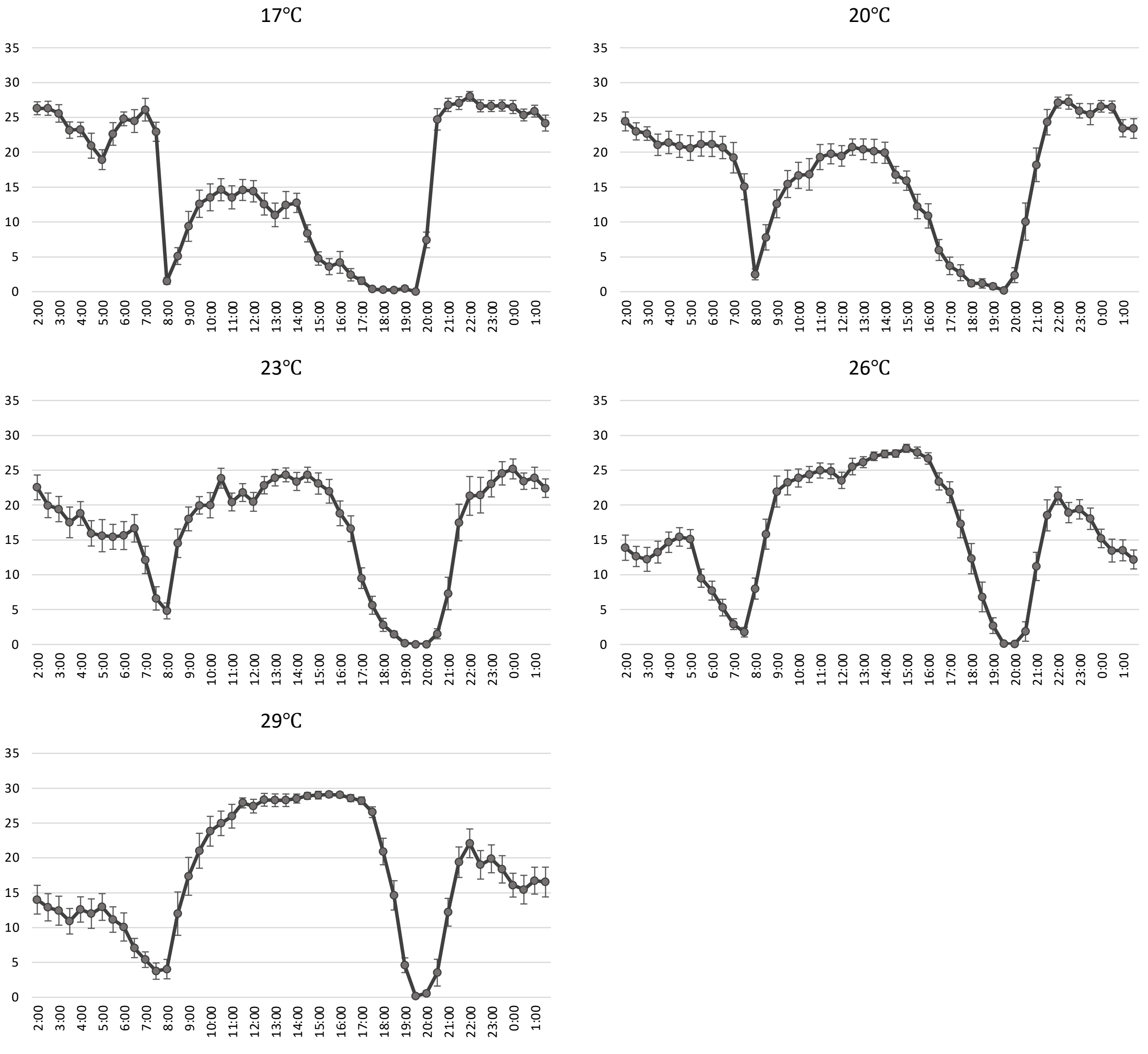


*D. melanogaster*

#### Supplementary Figure 7. (b)

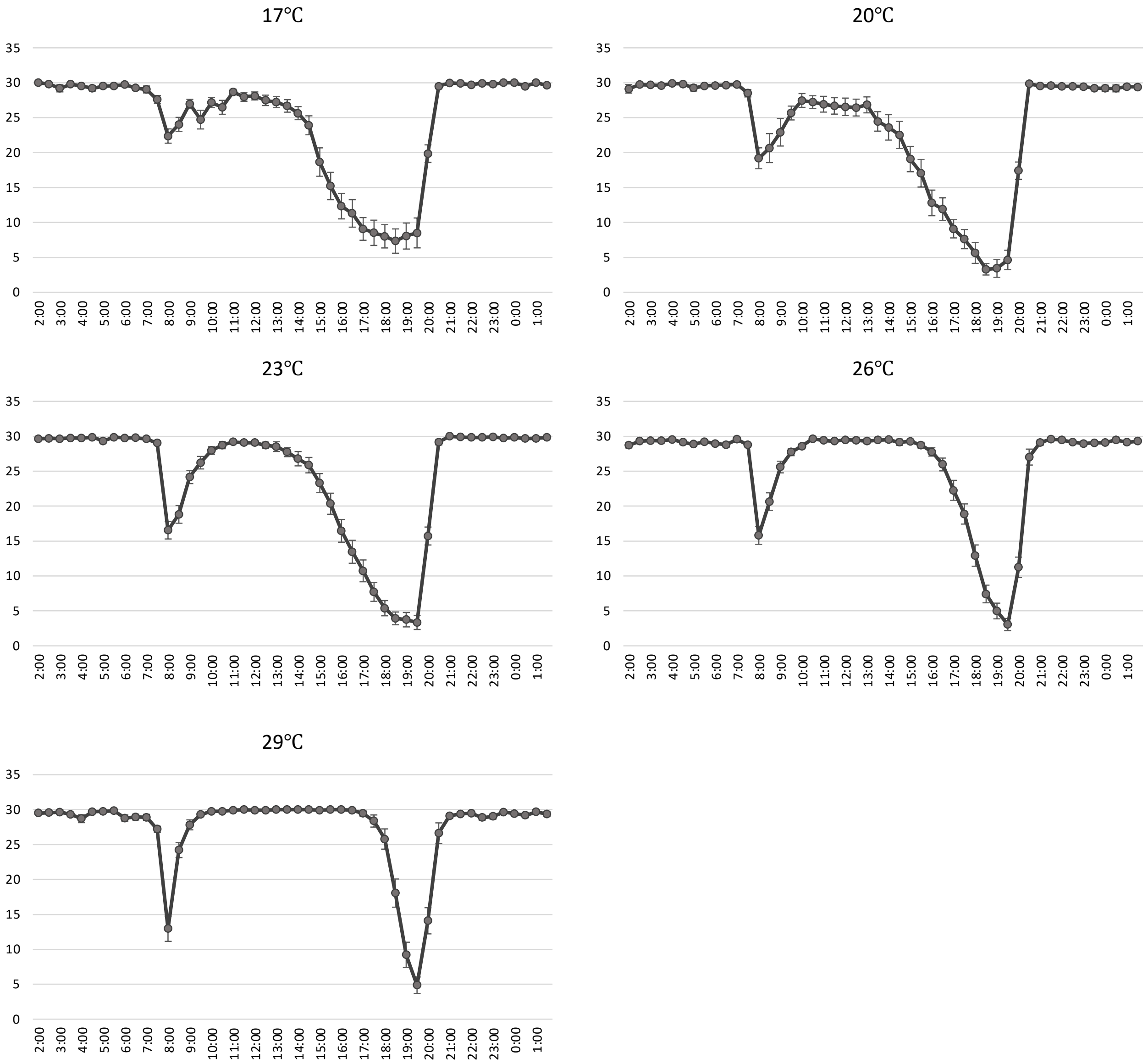


*D. simulans*

#### Supplementary Figure 7. (c)

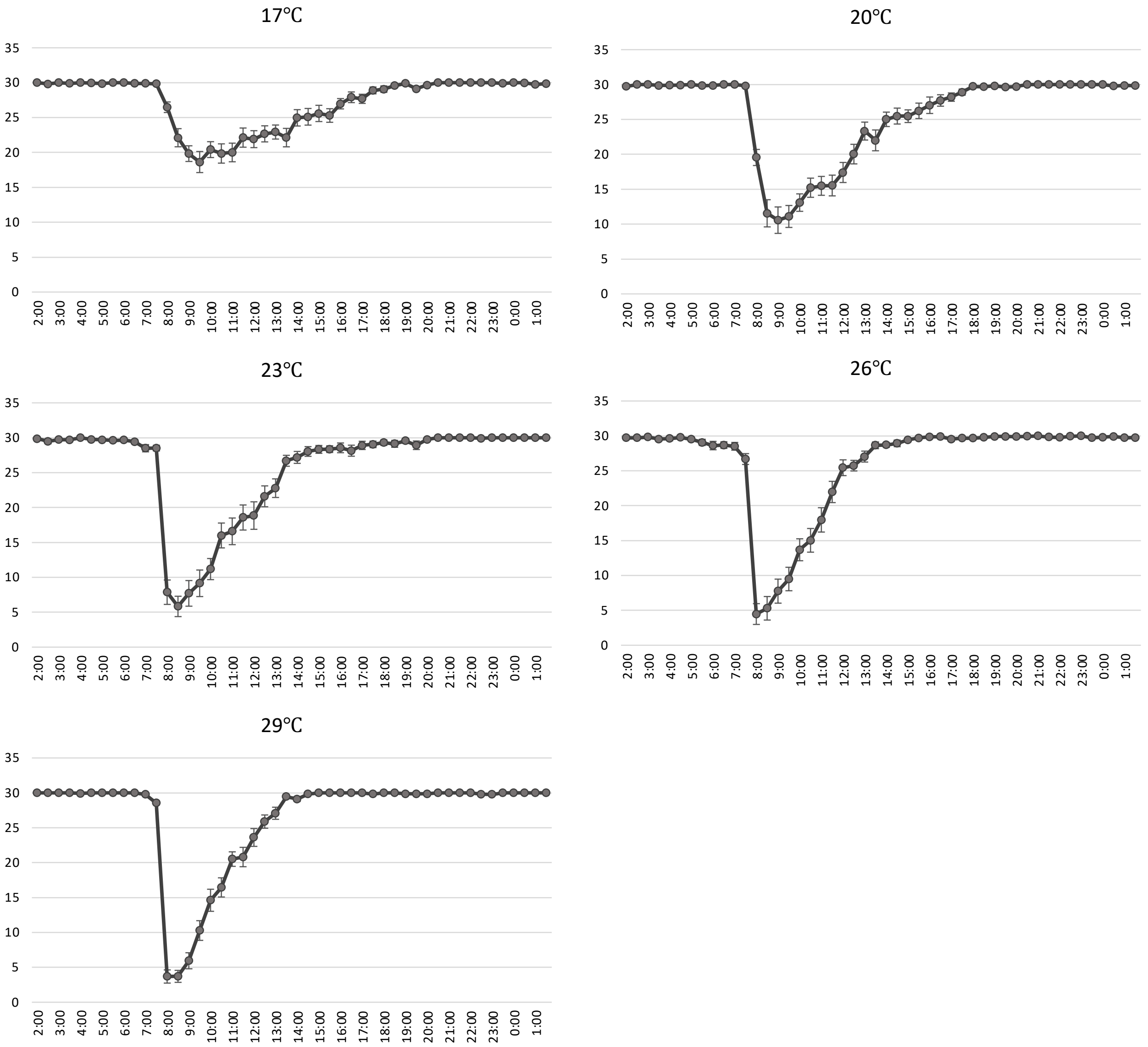


*D. ananassae*

Supplementary Figure 7. (d)

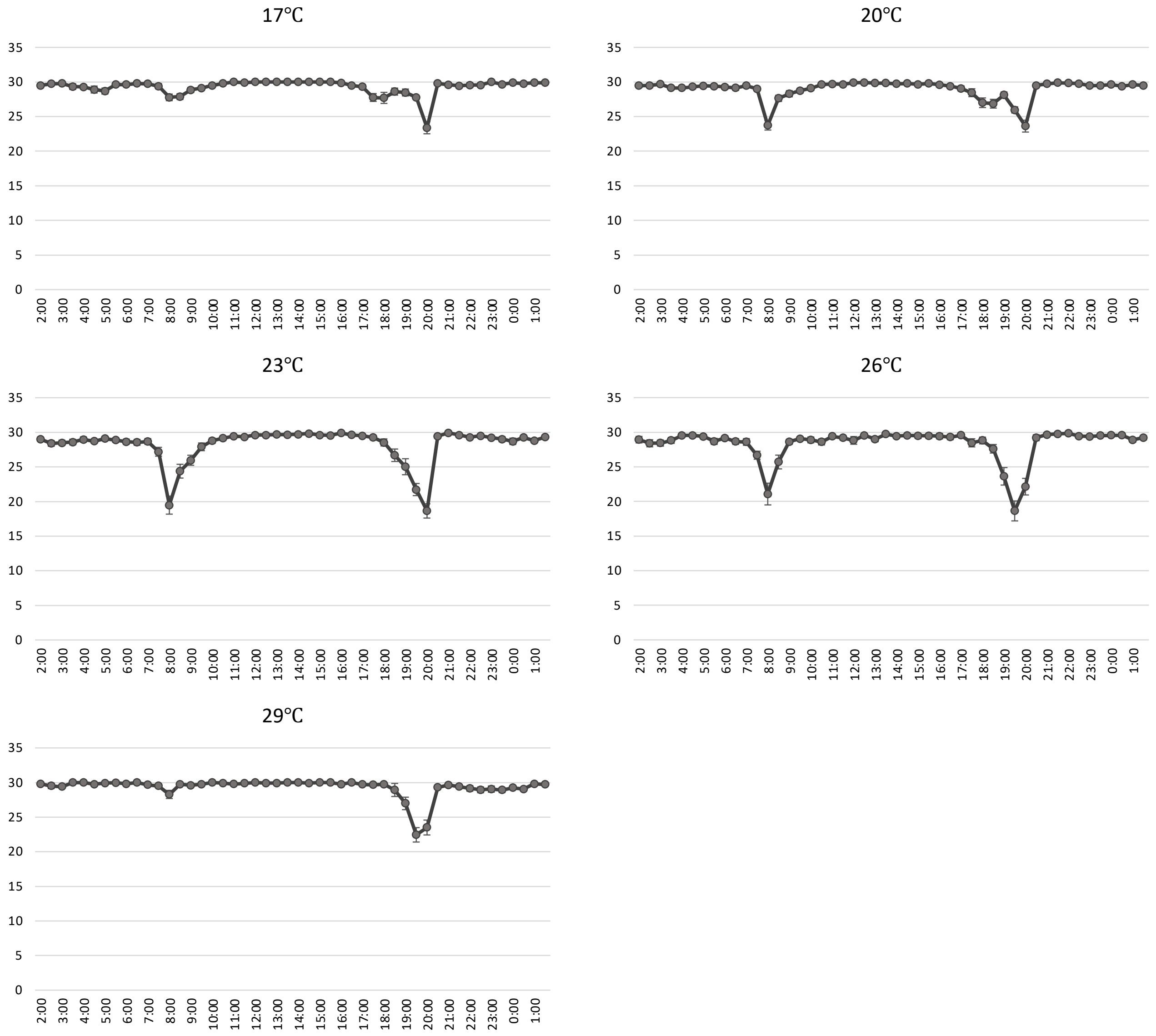


*D. erecta*

Supplementary Figure 7. (e)

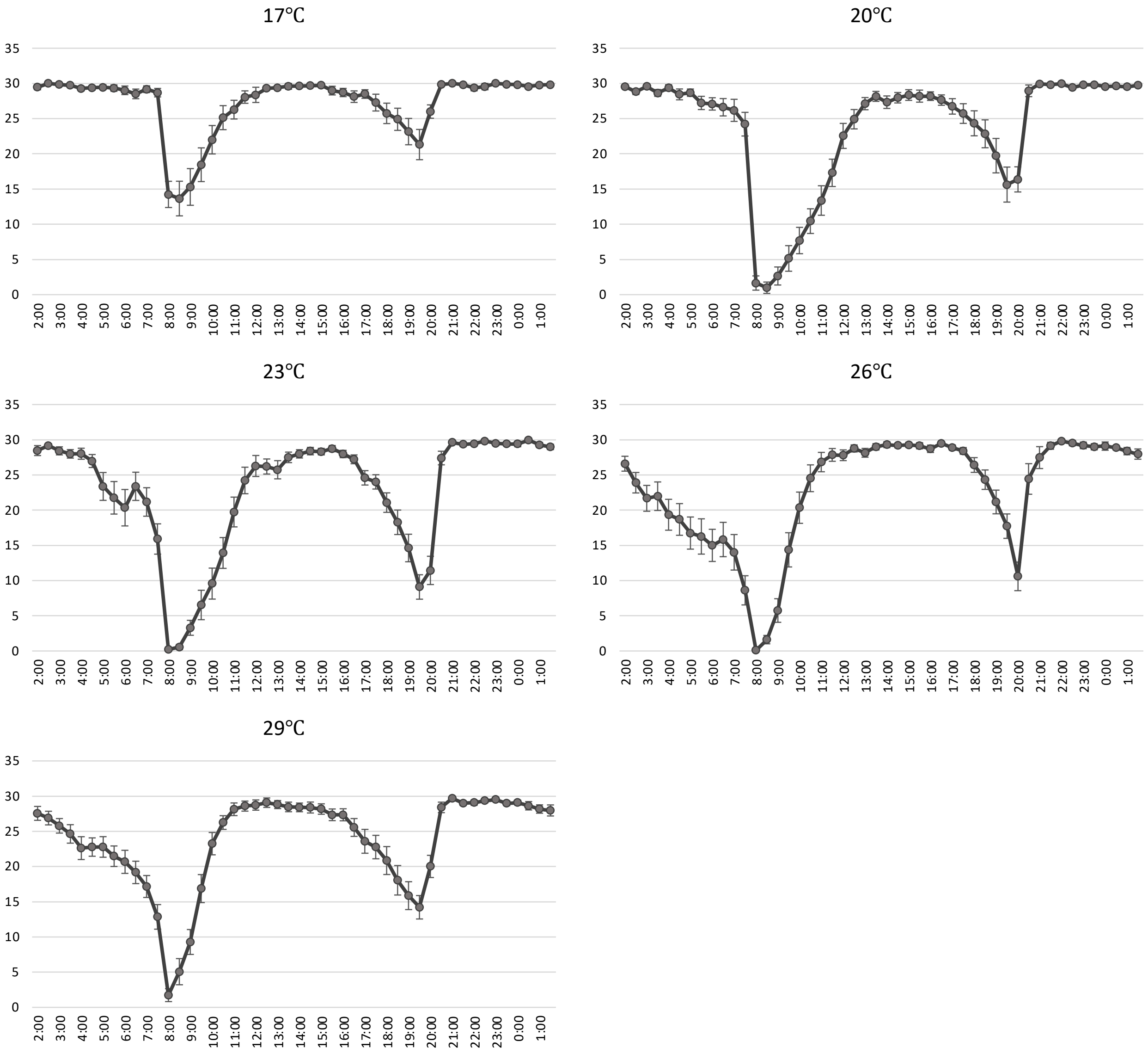


*D. yakuba*

Supplementary Figure 7. (f)

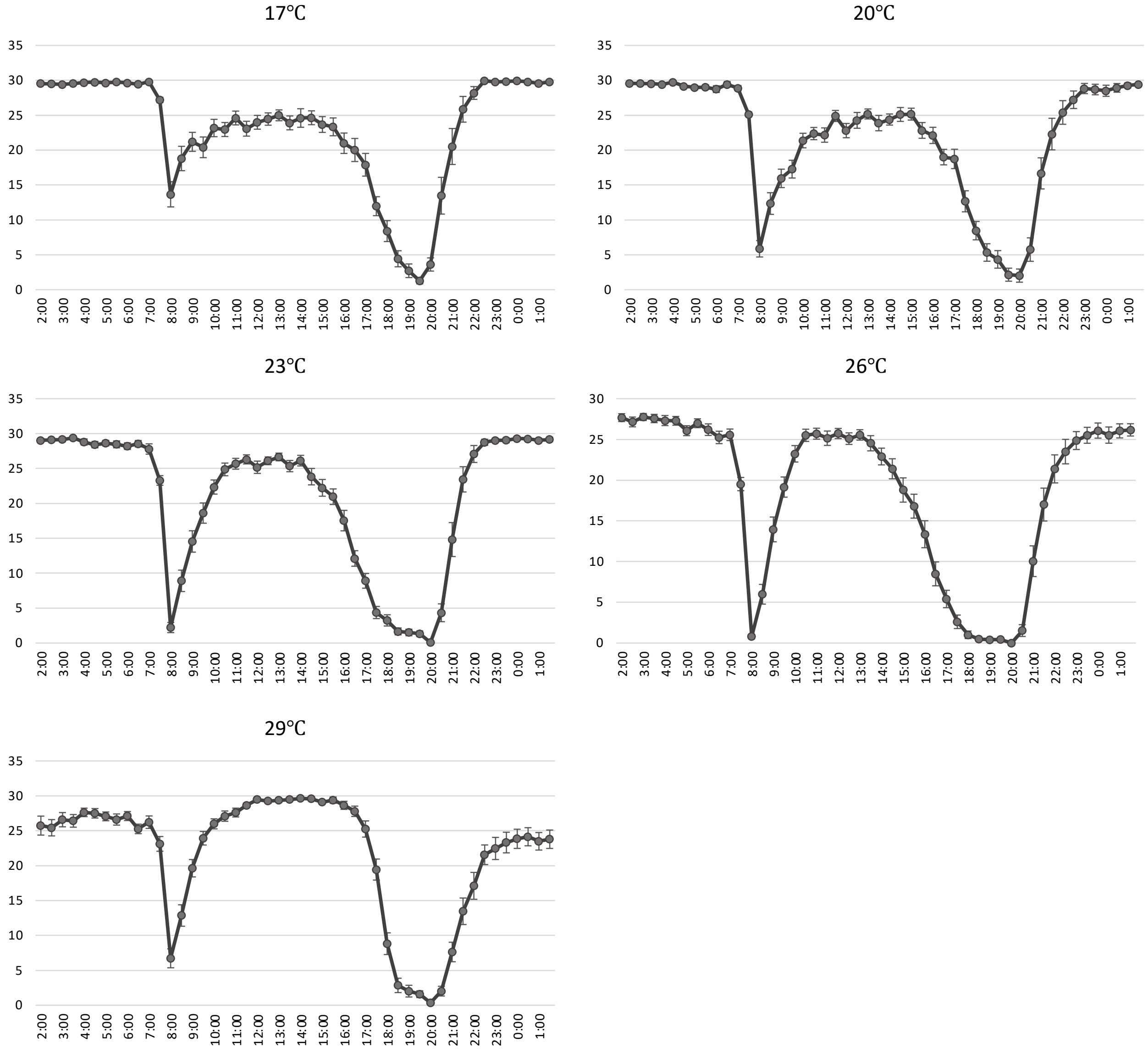


*D. sechellia*

Supplementary Figure 7. (g)

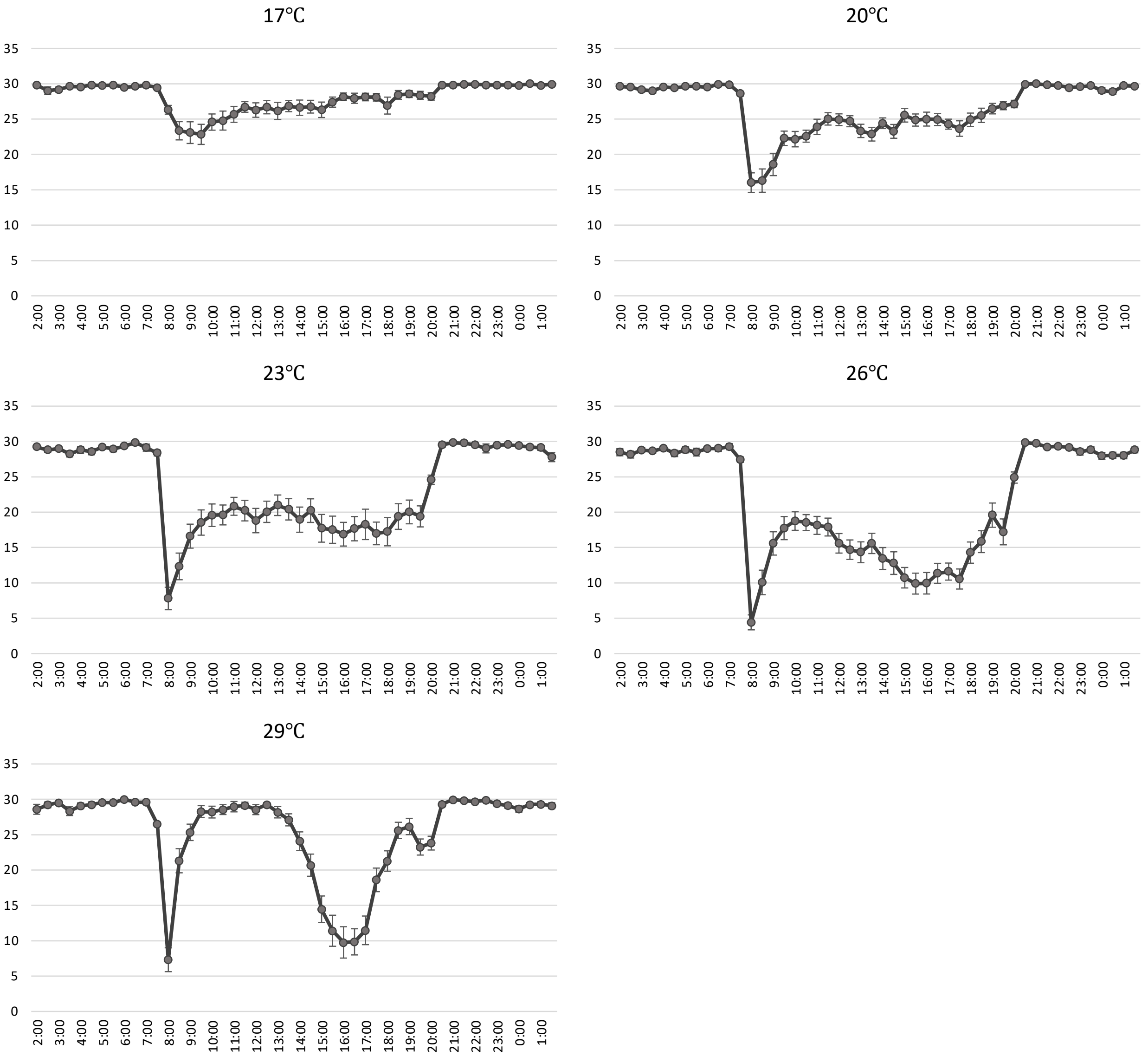


*D. willistoni*

Supplementary Figure 7. (h)

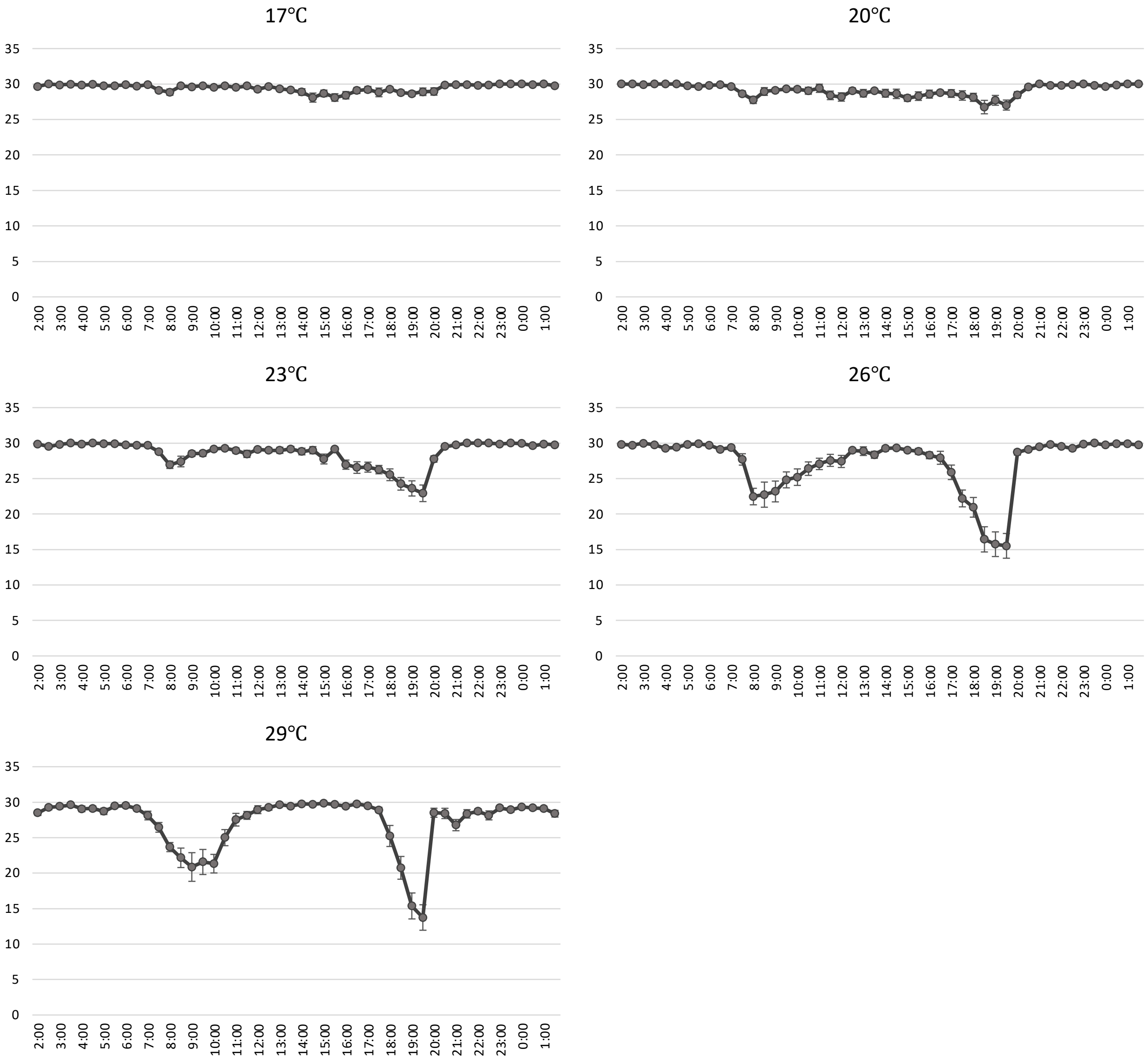


*D. mojavensis*

Supplementary Figure 7. (i)

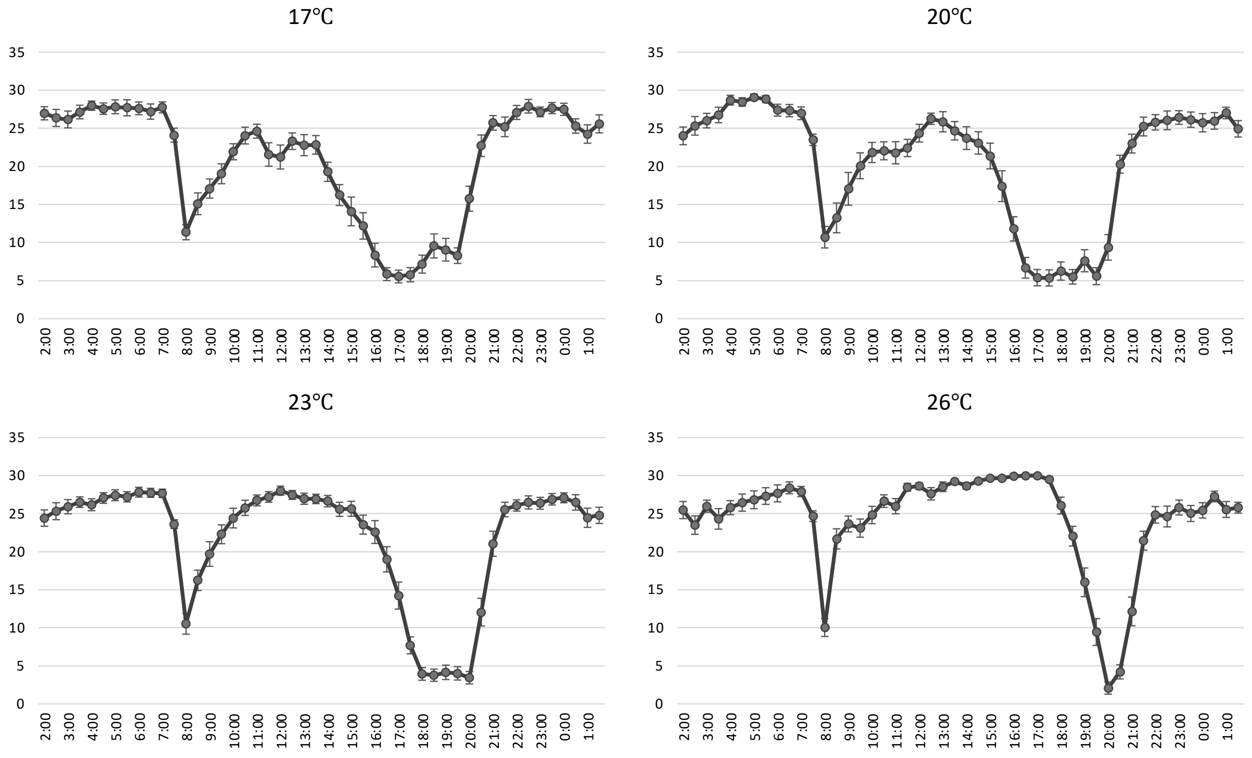


*D. persimilis*

Supplementary Figure 7. (j)

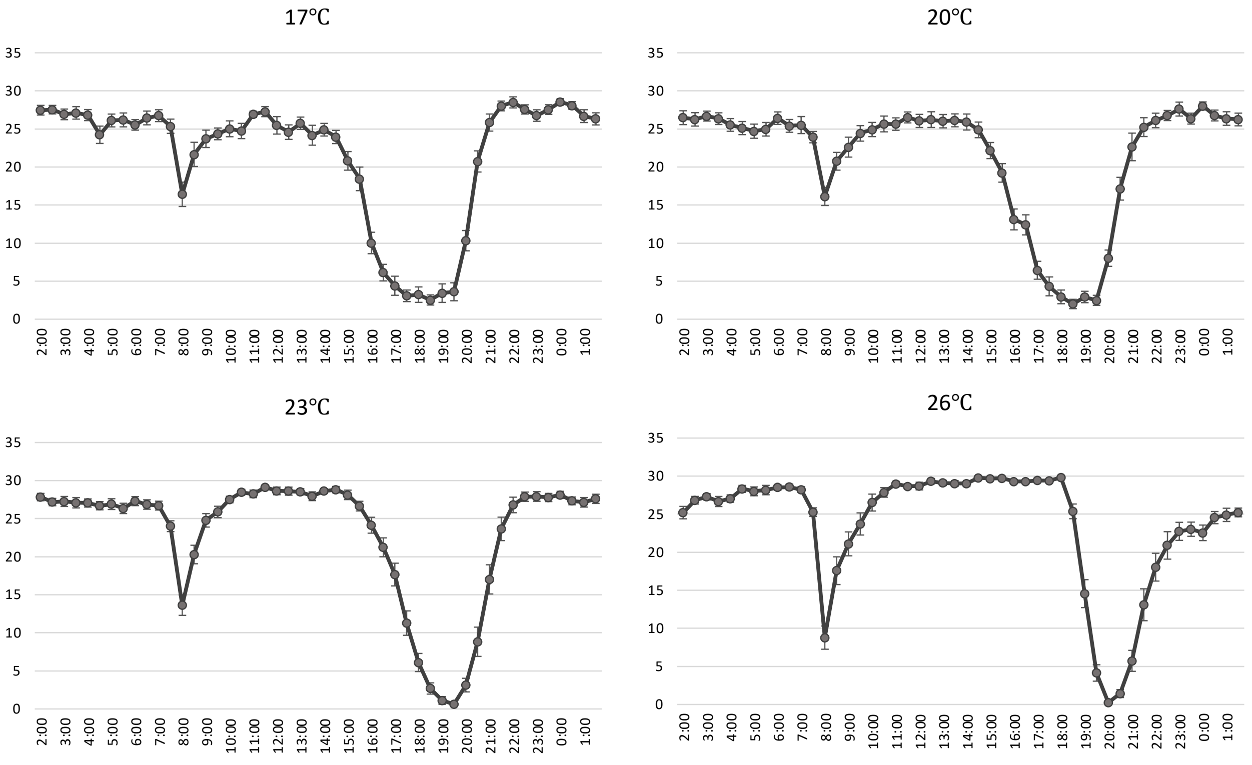


*D. pseudoobscura*

#### Supplementary Figure 7. (k)

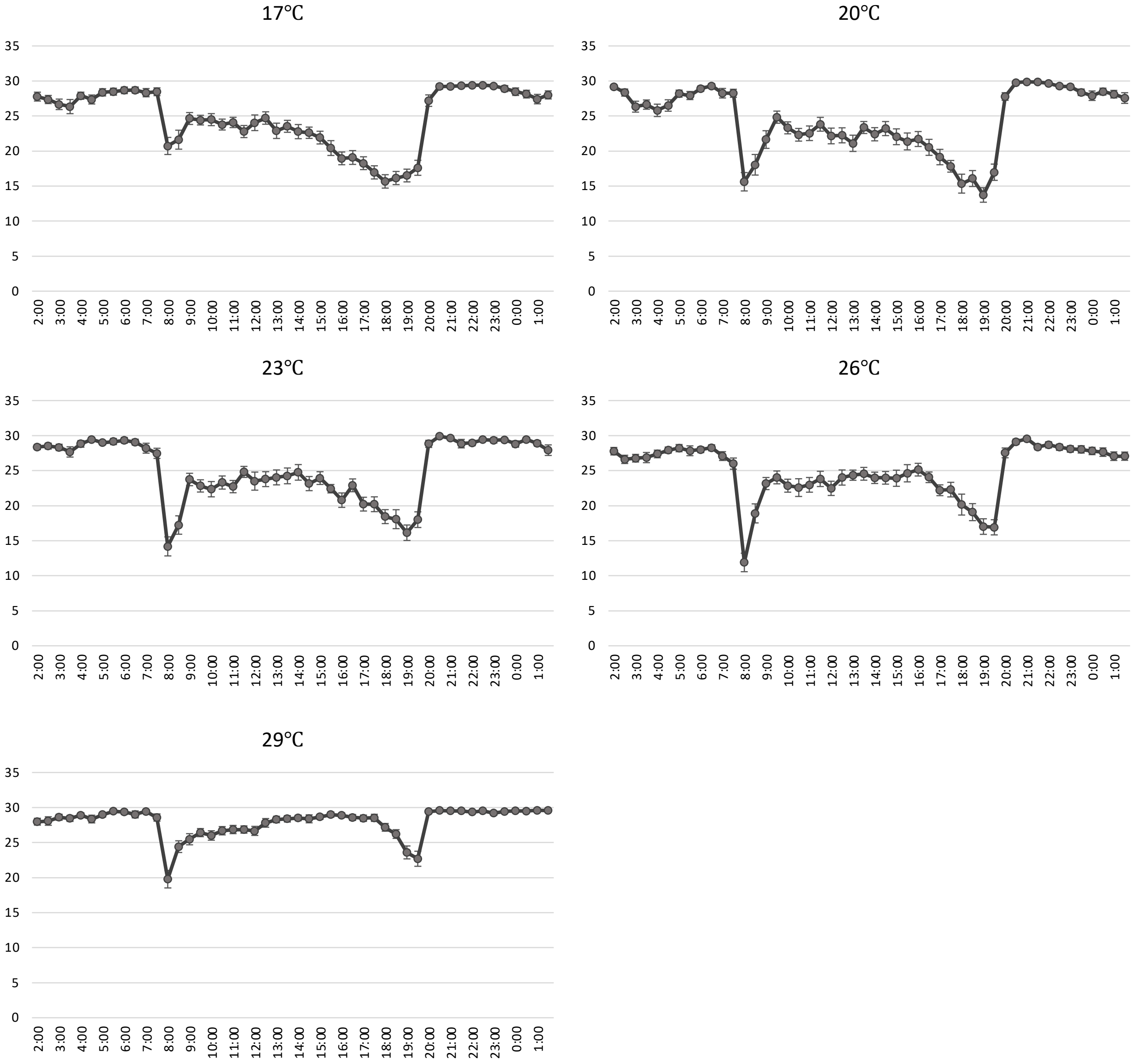


*D. virilis*

Supplementary Figure 8.

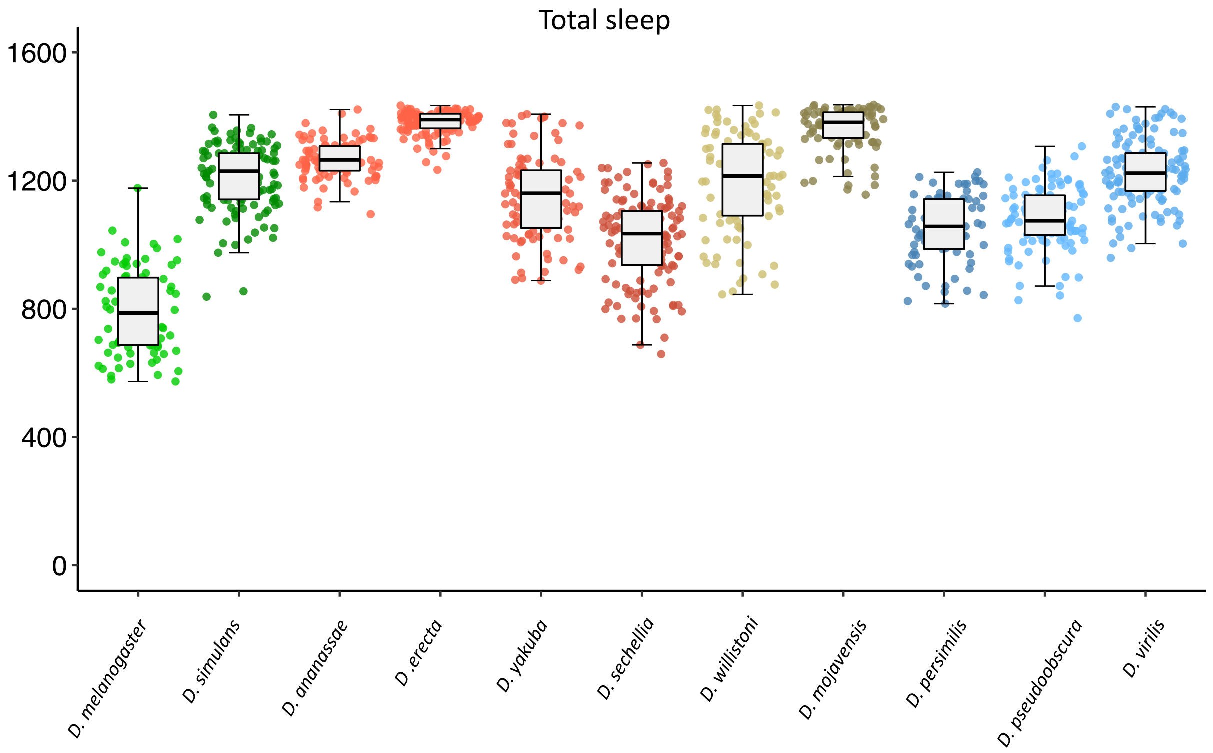

Supplementary Figure 9.

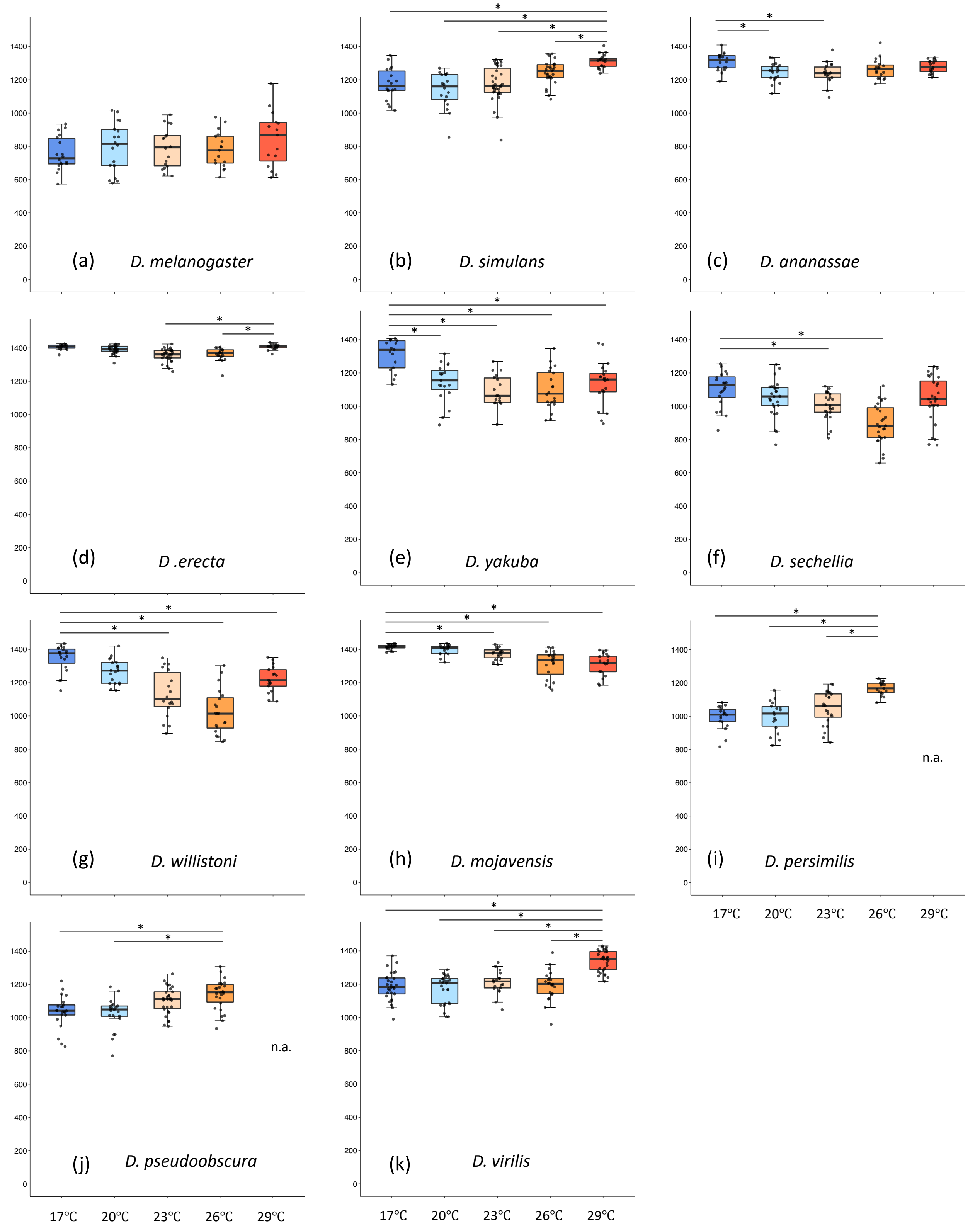

Supplementary Figure 10.

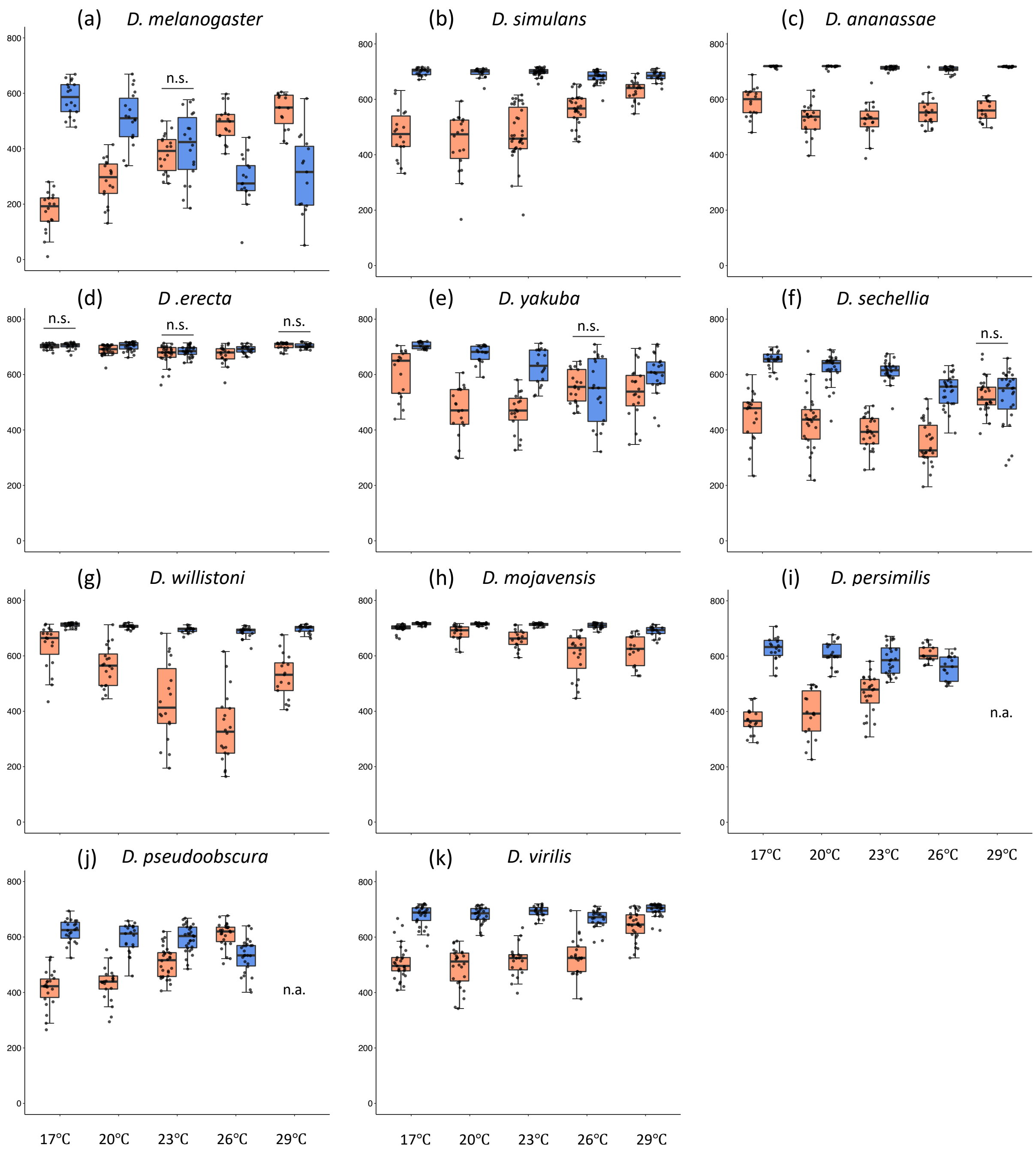

Supplementary Figure 11.

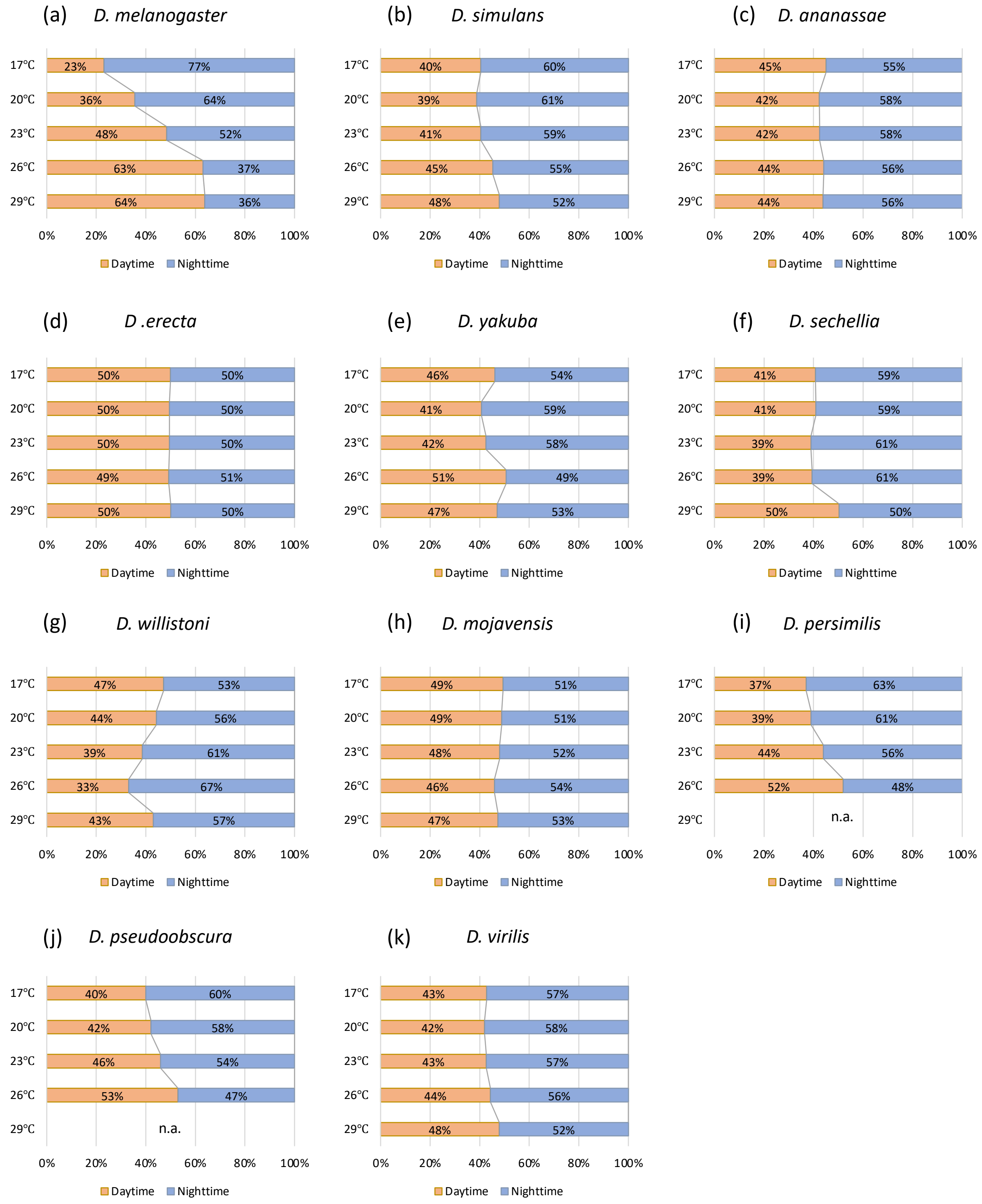

Supplementary Figure 12.

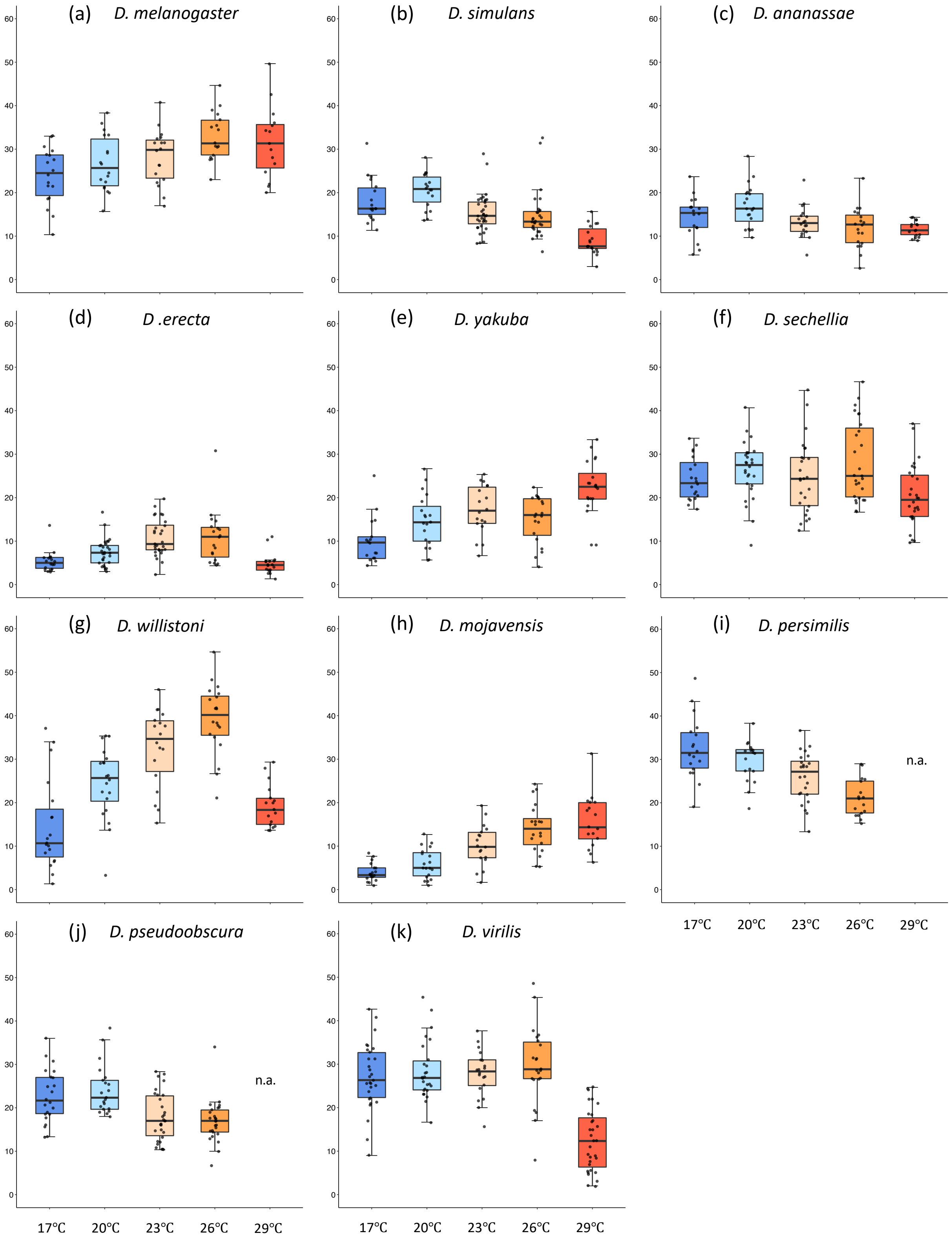

Supplementary Figure 13.

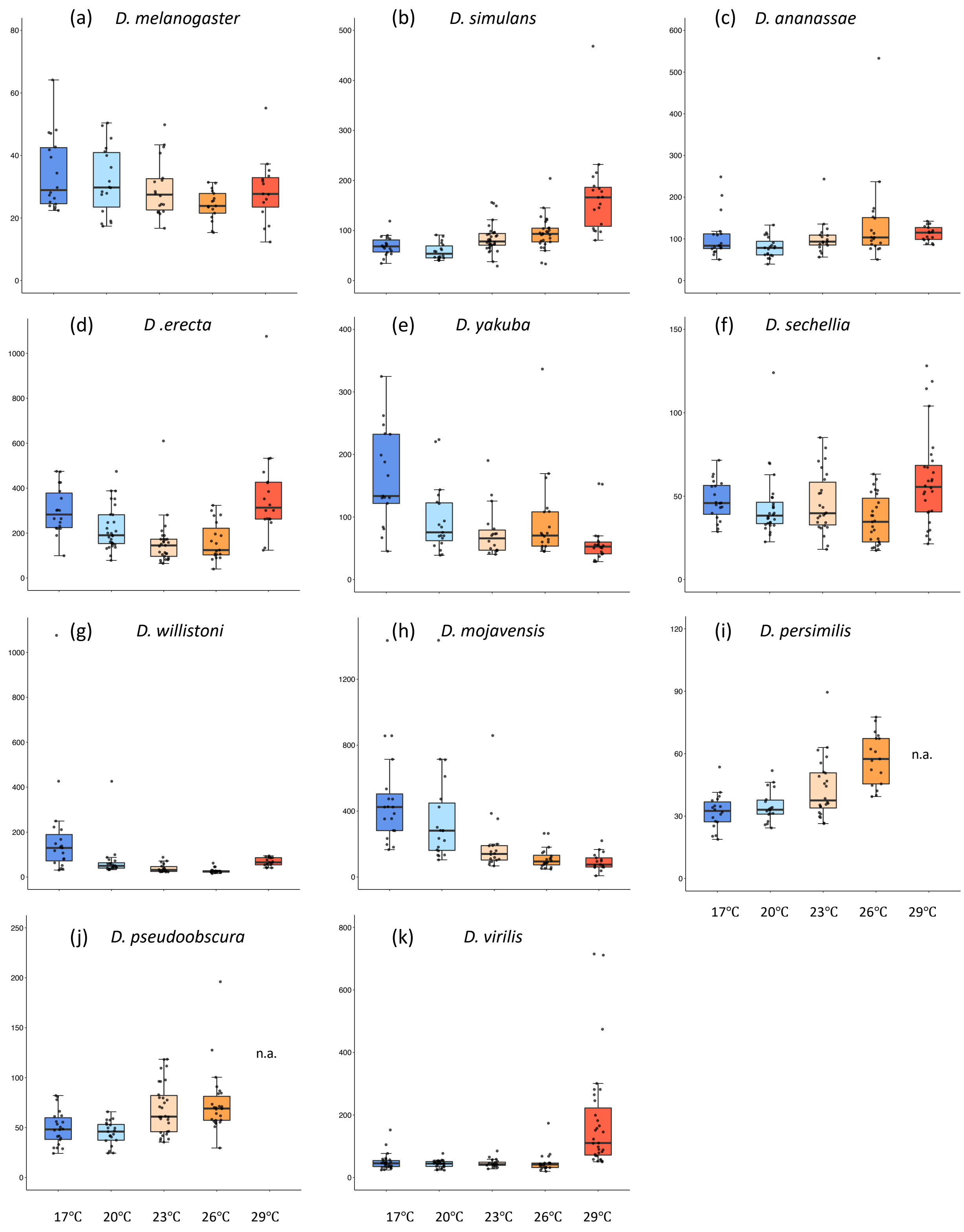

#### Supplementary Table 1.

| **Species** | *Total activity vs total sleep* | *Total sleep vs sleep number* | *Total sleep vs single sleep duration* |
|---|---|---|---|
| *D. melanogaster* | **-0.762** | -0.078 | **0.549** |
| *D. simulans* | **-0.871** | **-0.622** | **0.738** |
| *D. ananassae* | **-0.829** | **-0.412** | **0.509** |
| *D .erecta* | **-0.864** | **-0.963** | **0.969** |
| *D. yakuba* | **-0.908** | **-0.615** | **0.753** |
| *D. sechellia* | **-0.814** | **-0.550** | **0.749** |
| *D. willistoni* | **-0.958** | **-0.897** | **0.936** |
| *D. mojavensis* | **-0.843** | **-0.981** | **0.983** |
| *D. persimilis* | **-0.796** | **-0.674** | **0.837** |
| *D. pseudoobscura* | **-0.607** | **-0.673** | **0.787** |
| *D. virilis* | **-0.697** | **-0.903** | **0.936** |

Supplementary Table S1. Spearman rank correlations between total activity and total sleep, total sleep and sleep number, and total sleep and single sleep duration. The bold values denote significant correlations.

**Supplementary Figure Legends**

Supplementary Figure 1. Total daily locomotor activities of 11 *Drosophila* species. Each dot indicates average total daily activity (count) of individual fly. The median values are as follows; *D. melanogaster* (median = 829.3), *D. simulans* (median = 215.0) *D. ananassae* (median = 89.3), *D. erecta* (median = 35.3), *D. yakuba* (median = 311.0), *D. sechellia* (median = 456), *D. williston*i (median = 104.8), *D. mojavensis* (median = 44.7). *D. persimilis* (median = 438.3), *D. pseudoobscura* (median = 557.3), and *D. virilis* (median = 221.5).

Supplementary Figure 2. Thermal performance curves for daily locomotor activity of 11 *Drosophila* species. Best fitting model (model), degree of freedom (*df*), optimum temperature ($T_{opt}$), and maximum performance ($P_{max}$) of each species are as follows. (a) *D. melanogaster* [model = gaussian_1987, *df* = 83, $T_{op}$ = 24.61ºC, $P_{max}$ = 920.3], (b) *D. simulans* [model = gaussian_1987, *df* = 116, $T_{op}$ = 20.59, $P_{max}$ = 256.72], (c) *D. ananassae* [model = gaussian_1987, *df* = 89, $T_{op}$ = 24.48, $P_{max}$ = 139.95], (d) *D. erecta* [model = flinn_1991, *df* = 110, $T_{op}$ = 23.86, $P_{max}$ = 67.22], (e) *D. yakuba* [model = gaussian_1987, *df* = 91, $T_{op}$ = 24.78, $P_{max}$ = 515.22], (f) *D. sechellia* [model = flinn_1991, *df* = 126, $T_{op}$ = 26.51, $P_{max}$ = 644.15], (g) *D. willistoni* [model = flinn_1991, *df* = 91, $T_{op}$ = 25.51, $P_{max}$ = 204.34], (h) *D. mojavensis* [model = quadratic_2008, *df* =90, $T_{op}$ = 29, $P_{max}$ = 107.47], (i) *D. persimilis* [model = gaussian_1987, *df* = 74, $T_{op}$ = 20.13, $P_{max}$ = 539.38], (j) *D. pseudoobscura* [model = gaussian_1987, *df* =96, $T_{op}$ = 23.19, $P_{max}$ = 632.57], and (k) *D. virilis* [model = quadratic_2008, *df* = 125, $T_{op}$ = 17, $P_{max}$ = 345.69]. Blue lines and dots indicate estimated TPCs and total daily locomotor activity of individual fly, respectively.

Supplementary Figure 3. Actograms of 11 *Drosophila* species at five different temperatures. (a) ~ (k) represents the actograms of each species; (a) *D. melanogaster*, (b) *D. simulans*, (c) *D. ananassae*, (d) *D. erecta*, (e) *D. yakuba*, (f) *D. sechellia*, (g) *D. willistoni*, (h) *D. mojavensis*, (i) *D. persimilis*, (j) *D. pseudoobscura*, and (k) *D. virilis*. Each bar shows activity counts per 30 min. The white bar indicates daytime and gray bar indicates nighttime activities. Error bars indicate standard error of means.

Supplementary Figure 4. Total locomotor activity in the first and second half of a day. (a) ~ (k) represents total locomotor activity of the first (orange) and second (blue) half of a day in each species; (a) *D. melanogaster*, (b) *D. simulans*, (c) *D. ananassae*, (d) *D. erecta*, (e) *D. yakuba*, (f) *D. sechellia*, (g) *D. willistoni*, (h) *D. mojavensis*, (i) *D. persimilis*, (j) *D. pseudoobscura*, and (k) *D. virilis*. The horizonal bars with n.s. above the pairs of box (daytime and nighttime) indicate no significant differences (Mann–Whitney U tests, $p > 0.05$), and all the rest of pairs show significant differences (Mann–Whitney U tests, $p < 0.05$). There was no significant difference in the amount of activity in the first half of a day among experimental temperatures in *D. simulans*, *D. pseudoobscura* and *D. virilis* (Kruskal–Wallis test: $p > 0.05$). Otherwise, the amount of the activity in the first and second half is significantly affected by temperatures (Kruskal–Wallis tests, $p < 0.05$).

Supplementary Figure 5. Ratio of the activity level in the first and second half of a day. (a) ~ (k) represents the ratio of the activity level in the first (orange) and second (blue) half of a day in each species; (a) *D. melanogaster*, (b) *D. simulans*, (c) *D. ananassae*, (d) *D. erecta*, (e) *D. yakuba*, (f) *D. sechellia*, (g) *D. willistoni*, (h) *D. mojavensis*, (i) *D. persimilis*, (j) *D. pseudoobscura*, and (k) *D. virilis*. The numbers in the bar indicate the % of the first-half day activity and second-half day activity in a day.

Supplementary Figure 6. Ratio of the activity level in the daytime and nighttime. (a) ~ (k) represents the ratio of the activity level in daytime (orange) and nighttime (blue) in each species; (a) *D. melanogaster*, (b) *D. simulans*, (c) *D. ananassae*, (d) *D. erecta*, (e) *D. yakuba*, (f) *D. sechellia*, (g) *D. willistoni*, (h) *D. mojavensis*, (i) *D. persimilis*, (j) *D. pseudoobscura*, and (k) *D. virilis*. The numbers in the bar indicate the % of daytime activity and nighttime activity in a day.

Supplementary Figure 7. Sleep profiles of 11 *Drosophila* species at five different temperatures. (a) ~ (k) represents the profiles of daily sleep of each species; (a) *D. melanogaster*, (b) *D. simulans*, (c) *D. ananassae*, (d) *D. erecta*, (e) *D. yakuba*, (f) *D. sechellia*, (g) *D. willistoni*, (h) *D. mojavensis*, (i) *D. persimilis*, (j) *D. pseudoobscura*, and (k) *D. virilis*. The y axis shows the duration of sleep in 30 min. Error bars indicate standard error of means.

Supplementary Figure 8. Total daily sleep duration of 11 *Drosophila* species. Each dot indicates average total daily sleep duration of individual fly. The median values are as follows; *D. melanogaster* (median = 787), *D. simulans* (median = 1229.3) *D. ananassae* (median = 1264.7), *D. erecta* (median = 1390.0), *D. yakuba* (median = 1160.5), *D. sechellia* (median = 1034.7), *D. williston*i (median = 1214.3), *D. mojavensis* (median = 1381.7). *D. persimilis* (median = 1058.0), *D. pseudoobscura* (median = 1074.7), and *D. virilis* (median = 1223.2).

Supplementary Figure 9. Total daily sleep duration of 11 *Drosophila* species at five different temperatures. (a) ~ (k) represents the total daily sleep duration in each species; (a) *D. melanogaster*, (b) *D. simulans*, (c) *D. ananassae*, (d) *D. erecta*, (e) *D. yakuba*, (f) *D. sechellia*, (g) *D. willistoni*, (h) *D. mojavensis*, (i) *D. persimilis*, (j) *D. pseudoobscura*, and (k) *D. virilis*. Caution that the y axis range are different among species. The box color indicates experimental temperatures; dark blue:17°C, light blue: 20°C, light orange: 23°C, dark orange: 26°C, red: 29°C. Each dot indicates average total daily sleep of individual fly. Except for *D. melanogaster*, the other species show the significant difference in total sleep duration among different temperature conditions (Kruskal–Wallis tests, $p < 0.05$). The horizonal bar with * above the boxes indicates the significant differences in the pairwise-comparison between the highest average of total sleep duration and others (Bonferroni/Dunn test, $p < 0.05$).

Supplementary Figure 10. Daytime and nighttime sleep of 11 *Drosophila* species at five different temperatures. (a) ~ (k) represents the daytime (orange) and nighttime (blue) sleep in each species; (a) *D. melanogaster*, (b) *D. simulans*, (c) *D. ananassae*, (d) *D. erecta*, (e) *D. yakuba*, (f) *D. sechellia*, (g) *D. willistoni*, (h) *D. mojavensis*, (i) *D. persimilis*, (j) *D. pseudoobscura*, and (k) *D. virilis*. The horizonal bars with n.s. above the box daytime and nighttime pairs indicate not significant differences (Mann–Whitney U tests, $p > 0.05$), otherwise each pairs shows a significant difference (Mann–Whitney U tests, $p < 0.05$).

Supplementary Figure 11. Ratio of daytime and nighttime sleep. (a) ~ (k) represents the ratio of daytime (orange) and nighttime (blue) sleep in each species; (a) *D. melanogaster*, (b) *D. simulans*, (c) *D. ananassae*, (d) *D. erecta*, (e) *D. yakuba*, (f) *D. sechellia*, (g) *D. willistoni*, (h) *D. mojavensis*, (i) *D. persimilis*, (j) *D. pseudoobscura*, and (k) *D. virilis*. The numbers in the bar indicate the % of the daytime sleep and nighttime sleep in total daily sleep.

Supplementary Figure 12. The number of daily sleep of 11 *Drosophila* species at five different temperatures. (a) ~ (k) represents the number of daily sleep in each species; (a) *D. melanogaster*, (b) *D. simulans*, (c) *D. ananassae*, (d) *D. erecta*, (e) *D. yakuba*, (f) *D. sechellia*, (g) *D. willistoni*, (h) *D. mojavensis*, (i) *D. persimilis*, (j) *D. pseudoobscura*, and (k) *D. virilis*. The box color indicates experimental temperatures; dark blue:17°C, light blue: 20°C, light orange: 23°C, dark orange: 26°C, red: 29°C. Each dot indicates average number of daily sleep of individual fly. In all species, the number of daily sleep is significantly affected by temperatures (Kruskal–Wallis tests, $p < 0.05$).

Supplementary Figure 13. Length of single sleep of 11 *Drosophila* species at five different temperatures. (a) ~ (k) represents the single sleep duration in each species; (a) *D. melanogaster*, (b) *D. simulans*, (c) *D. ananassae*, (d) *D. erecta*, (e) *D. yakuba*, (f) *D. sechellia*, (g) *D. willistoni*, (h) *D. mojavensis*, (i) *D. persimilis*, (j) *D. pseudoobscura*, and (k) *D. virilis*. The box color indicates experimental temperatures; dark blue:17°C, light blue: 20°C, light orange: 23°C, dark orange: 26°C, red: 29°C. Each dot indicates average single sleep duration of individual fly. In all species,

length of single sleep is significantly affected by temperatures (Kruskal–Wallis tests, $p < 0.05$)
Each.
